# Cerebellar induced VTA plasticity underlies chronic stress-induced depression-like behaviors

**DOI:** 10.64898/2026.05.06.723377

**Authors:** Seulgi Kang, Taehyeong Kim, Daun Kim, Soo-Ji Baek, Thomas J. McHugh, Yukio Yamamoto, Keiko Tanaka-Yamamoto

**Affiliations:** Brain Science Institute, Korea Institute of Science and Technology (KIST), Seoul 02792, Republic of Korea; Division of Bio-Medical Science and Technology, KIST School, Korea University of Science and Technology (UST), Seoul 02792, Republic of Korea; Department of Integrated Biomedical and Life Sciences, Korea University, Seoul 02708, Republic of Korea; Department of Life Sciences, Korea University, Seoul 02841, Republic of Korea; Laboratory for Circuit and Behavioral Physiology, RIKEN Center for Brain Science, Wako-shi, Saitama 351-0198,Japan

## Abstract

Synaptic regulation is a key mechanism underlying neural circuit homeostasis and plasticity. We previously showed that projections from the deep cerebellar nuclei (DCN) to the ventral tegmental area (VTAp–DCN neurons) contribute to depression-like behaviors following chronic restraint stress (RS), but the mechanisms by which these outputs induce downstream long-term functional changes remain unknown. Here, we show that chronic RS induces an activity-dependent reduction in vesicular glutamate transporter 2 (VGLUT2) expression in VTAp–DCN axons, resulting in decreased miniature excitatory postsynaptic current frequency in dorsolateral VTA dopaminergic neurons and reduced phasic dopamine release in the nucleus accumbens. Notably, VGLUT2 reduction temporally coincides with the emergence of depression-like behaviors. Consistent with this, targeted VGLUT2 overexpression in VTAp–DCN neurons rescues synaptic transmission and dopamine signaling and prevents depression-like behaviors following chronic RS. These findings reveal a stress-induced plasticity mechanism in the cerebellar–VTA circuit linking chronic stress to depression-like behaviors.

## Introduction

The cerebellum can transmit information to influence distal targets via its efferent projections, primarily from the deep cerebellar nuclei (DCN) composed of the fastigial, interposed, and dentate (DN) nuclei ^1–3^. Recent advances in pathway-specific manipulation techniques have enabled the dissection of the functions of individual DCN projections, deepening our understanding of the distinct roles the structure can play ^4^. As a result, the traditional view of the cerebellum as a motor structure has been expanded to include its role in a range of non-motor functions, including cognitive and affective processes ^5–14^. These emerging functions appear to be mediated by specific DCN efferent neurons modulating the activity of the responsible target brain regions. However, how the cerebellum contributes to downstream activity-dependent long-lasting changes, i.e., plasticity, in response to environmental or physiological conditions remains largely unknown.

One of the brain regions innervated by DCN neurons is the ventral tegmental area (VTA) ^6,10,15^. The VTA plays a central role in reward and motivation by regulating dopamine (DA) release ^16–18^. It is also critically involved in stress responses, as stress exposure alters VTA neuronal activity, synaptic inputs, and molecular expression profiles ^19–22^. The resulting stress-induced changes in DA release across VTA projection targets are thought to shape emotional and behavioral responses ^23–26^. However, the nature of these changes varies considerably depending on factors such as the modality and duration of the stress, the location and timing of DA measurement, and the specific afferent pathways engaged under each condition ^20,22,27–29^. Electrical or chemogenetic stimulation of the DN has been reported to trigger DA release in the nucleus accumbens (NAc) ^30,31^, a major target of VTA dopaminergic (DAergic) neurons, suggesting that the cerebellar afferents may contribute to stress-related changes in VTA function through synaptic or molecular modulation.

We previously demonstrated that DCN neurons projecting to the VTA (VTAp–DCN neurons) regulate the development of depression-like behaviors induced by chronic restraint stress (RS) ^10^. Specifically, both short-term (3 days) and chronic (14 days) stress activate DCN neurons. Moreover, chemogenetic inhibition of VTAp–DCN neurons during chronic RS prevents the emergence of depression-like behaviors, whereas chronic excitation of these neurons alone is sufficient to induce a comparable behavioral phenotype even in the absence of RS. Given that depression-like behaviors are generally attributed to prolonged rather than brief stress exposure, we hypothesized that repeated activation of VTAp–DCN neurons by chronic RS triggers plastic changes in their downstream targets, which in turn lead to the behavioral phenotype. Here, we show that RS induces a delayed and persistent reduction in vesicular glutamate transporter 2 (VGLUT2) expression in VTAp–DCN neurons via activity-dependent mechanisms. This reduction exerts a causal influence on depression-like behaviors through modulation of DA release, thereby elucidating a molecular and synaptic mechanism by which the cerebellum contributes to stress-induced depression.

## Results

### Depression-like behaviors triggered by 14RS, but not 3RS

We first characterized depression-like behaviors in mice exposed to acute or chronic stress. Mice were subjected to RS for 2 h daily over either 3 or 14 consecutive days, referred to as 3RS and 14RS, respectively, with the latter representing a chronic RS paradigm (Figure 1A). Blood corticosterone levels increased immediately after both 3RS and 14RS (Figure 1B), confirming that RS effectively induced stress in mice, though this elevation was no longer observed 10 days after 14RS (14RS+10, Figure 1B).

**Figure 1.**
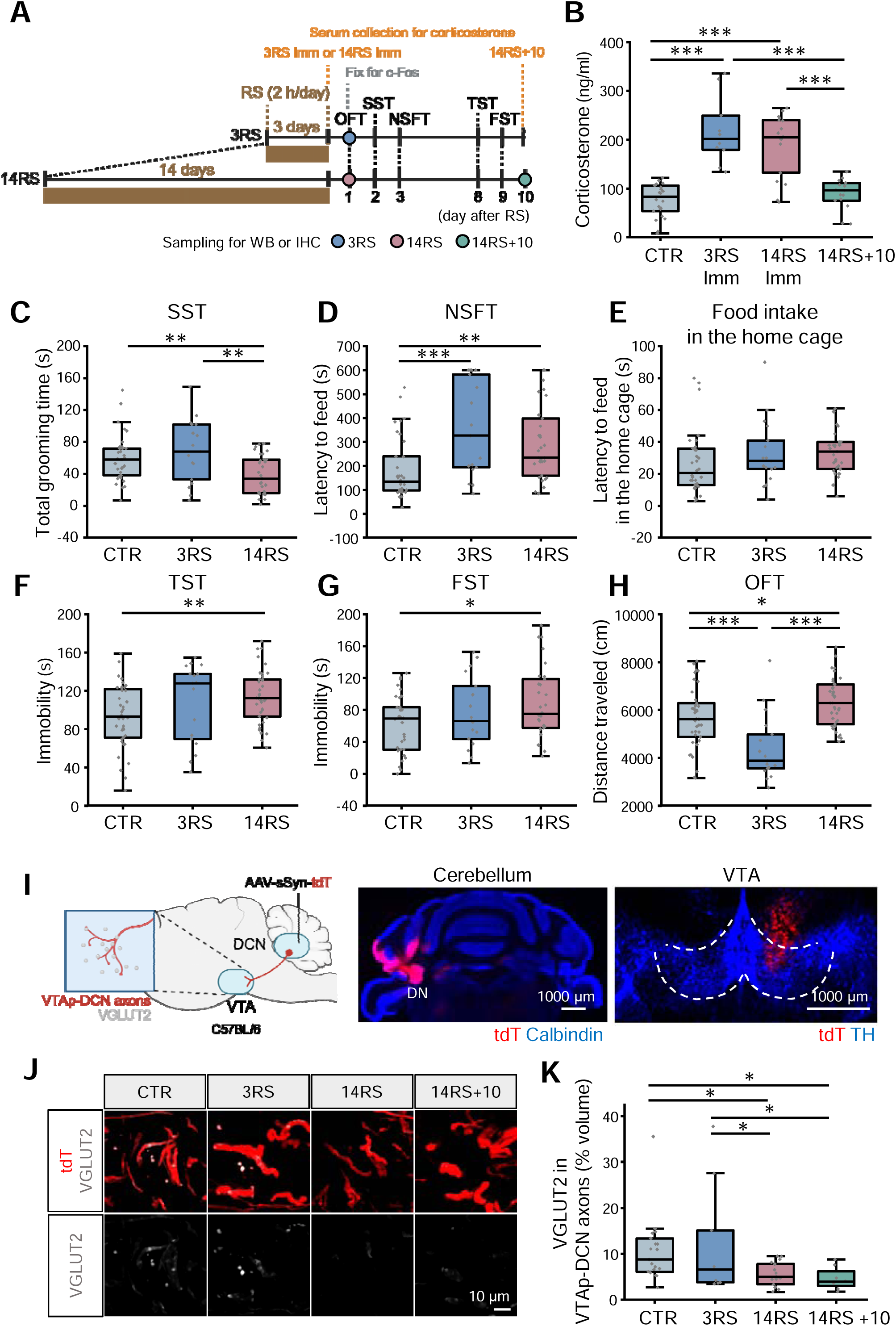
Temporal coupling of depression-like behaviors and VGLUT2 reduction in VTAp-DCN axons. (A) Timeline of behavioral tests, sample collection (colored circles) for western blot (WB) and immunohistochemistry (IHC) of synaptic molecules, corticosterone measurements, and fixation for c-Fos immunostaining. 3RS or 14RS indicates analyses performed or initiated one day after RS cessation, whereas 14RS+10 indicates analyses performed at 10 days after the end of 14RS. Note that corticosterone was measured immediately after RS (3RS Imm or 14RS Imm). (B) Serum corticosterone levels in CTR (n = 22 mice), 3RS Imm (n = 10 mice), 14RS Imm (n = 18 mice), and 14RS+10 (n = 15 mice) groups. (C–H) Total grooming time in the SST (C), latency to feed in the NSFT (D) and in the home cage (E), immobility time in the TST (F) and FST (G), and total distance traveled in the OFT (H), comparing 3RS (n = 17 mice) and 14RS (n = 31 in (C–E); n = 38 in (F); n = 30 in (G); n = 39 in H) groups with CTR (n = 34 in (C–E) and (G); n = 38 in (F) and (H)). (I) Schematic of DCN neuron labeling via AAV injection and representative images of tdT expression (red) in a cerebellar slice stained with calbindin (blue) and a midbrain slice stained with TH (blue). (J) Representative 3D projection images of tdT-labeled VTAp–DCN axons (red) and VGLUT2 staining (gray) restricted to tdT-positive VTAp–DCN axons. (K) Percentage of VGLUT2-positive volume within VTAp–DCN axons in CTR (n = 19 mice), 3RS (n = 9 mice), 14RS (n = 17 mice), and 14RS+10 (n = 8 mice). *p < 0.05, **p < 0.01, ***p < 0.001; one-way ANOVA followed by Fisher’s LSD post hoc test. All data in this and subsequent figures are presented as boxplots with gray dots representing individual data points, center lines denoting the median, boxes representing the 25th–75th percentiles, and whiskers indicating the most extreme values excluding outliers. Exact p-values for the datasets in this and subsequent figures are provided in Table S1.

To assess depression-like behaviors, we conducted the sucrose splash test (SST), novelty-suppressed feeding test (NSFT), tail suspension test (TST), and forced swimming test (FST) spaced across the 10 days following the end of RS (Figure 1A). As expected ^10,32–35^, mice in the 14RS group exhibited clear depression-like behaviors, including reduced grooming time in the SST (Figure 1C), increased latency to feed in the NSFT (Figure 1D) with no changes in home cage food consumption (Figure 1E), and increased immobility in both the TST (Figure 1F) and FST (Figure 1G) compared with the unstressed control (CTR) mice. Because travel distance in the open field test (OFT) was increased in the 14RS group (Figure 1H), the increased immobility observed in the TST and FST is unlikely to reflect reduced general locomotor activity. In contrast, the 3RS group did not exhibit clear depression-like behaviors in the SST, TST, or FST (Figures 1C, 1F and 1G), although these mice showed increased latency to feed in the NSFT (Figure 1D) and reduced travel distance in the OFT (Figure 1H), possibly reflecting anxiety-like responses ^36,37^. Notably, depression-like behaviors persisted for up to 10 days after the end of 14RS, as observed in the FST conducted 9 days post-stress (Figure 1G) and in our previous study ^10^. This contrasts with DCN neuronal activity assessed by c-Fos expression, which, although elevated immediately after both 3RS and 14RS ^10^, returned to control levels one day after 14RS (CTR, 7.4 ± 2.5%; 14RS, 4.3 ± 1.2%; n = 8 images from 4 mice per group; *p* = 0.43, Mann–Whitney U test). These findings indicate a temporal dissociation between DCN neuronal activation and the expression of depression-like behaviors, suggesting that although increased DCN neuronal activity triggers depression-like behaviors in response to chronic RS ^10^, it is not directly involved in their manifestation once these behaviors emerge, but instead initiates downstream changes that underlie delayed and persistent behavioral outcomes.

### Cerebellum-dependent plastic changes in synaptic molecules in the VTA

To identify such downstream mechanisms, we examined RS-dependent changes in excitatory synaptic molecules in the dorsolateral VTA, the primary target of DCN projections ^10^. We analyzed samples one day after 3RS or 14RS, as well as at 14RS+10 (Figure 1A, colored circles), allowing us to evaluate whether molecular changes temporally coincided with the emergence of depression-like behaviors. First, we examined several synaptic molecules using western blot analysis in VTA lysates. AMPA-type glutamate receptor GluR2 was significantly increased after both 3RS and 14RS, but not at 14RS+10, whereas postsynaptic density protein 95 (PSD-95) was significantly increased only at 14RS+10, but not after 3RS or 14RS, compared with CTR mice Figures S1B and S1C). Immunohistochemical staining for GluR2 and tyrosine hydroxylase (TH) in mice injected with AAV expressing tdTomato (tdT) in the DCN confirmed increased receptor expression in the VTA at the site of DCN innervation (Figures S1D–S1G). Consistent with the western blot results, GluR2 levels in the dorsolateral VTA were elevated after both 3RS and 14RS, but returned to baseline by 10 days after 14RS (Figures S1D and S1E). When GluR2 expression was examined specifically in TH-positive DAergic neurons, we found it was also elevated after 14RS but was not sustained at 14RS+10 (Figures S1F and S1G). However, as the temporal profiles of GluR2 and PSD-95 changes did not align with the time course of depression-like behaviors, these alterations are unlikely to be directly responsible for their manifestation, although they may be partially involved in the behaviors.

Stress-dependent depression-like behaviors have been associated with alterations in glutamatergic synapses, as indicated by changes in VGLUT2 expression ^38–40^. Despite no change in total VGLUT2 levels detected by western blot analysis of VTA lysates (Figures S1B and S1C), given that VTAp–DCN neurons are glutamatergic ^6,10^, express VGLUT2, and mediate chronic RS-induced depression-like behaviors via their activity, we next asked if RS induces VGLUT2 alterations specifically at DCN axon terminals. To address this, we analyzed VGLUT2 immunohistochemical signals in the dorsolateral VTA of mice injected with an AAV expressing tdT into the DCN (Figure 1I), after 3RS, 14RS, or at 14RS+10 (Figure 1A). Consistent with the western blot results, overall VGLUT2 levels in the dorsolateral VTA remained unchanged at all time points (Figures S2A and S2B). In contrast, VGLUT2 signals colocalized with tdT-positive VTAp–DCN axons were significantly reduced one and ten days after 14RS, but not after 3RS, compared with CTR mice (Figures 1J and 1K), temporally coinciding with the emergence of depression-like behaviors. The total volume of tdT-positive VTAp–DCN axons was unchanged across groups (Figure S2C), indicating that VGLUT2 reduction was not attributable to altered axonal volume. In addition, fluorescent in situ hybridization revealed no differences in VGLUT2 mRNA (Slc17a6) expression in the DN—the primary nucleus within the DCN projecting to the VTA ^10^—between CTR and 14RS mice (Figures S2D and S2E), suggesting regulation at the protein level. Together, these results indicate that chronic, but not short-term, RS induces a sustained reduction in VGLUT2 in VTAp–DCN neurons during the period when depression-like behaviors are expressed. This temporal correlation raises the possibility of a direct association between the VGLUT2 reduction in VTAp–DCN neurons and the manifestation of these behaviors.

We next examined whether the VGLUT2 reduction observed after 14RS depends on the activity of VTAp–DCN neurons. To this end, we employed an established rAAV2-retro chemogenetic strategy, previously used to demonstrate that depression-like behaviors depend on VTAp–DCN neuronal activity ^10^, to express the inhibitory chemogenetic receptor hM4Di (Gi) along with GFP specifically in VTAp–DCN neurons, and suppressed neuronal activity during RS by daily intraperitoneal injections of clozapine-N-oxide (CNO) (Figure 2A). In mice expressing GFP alone in VTAp–DCN neurons and receiving either CNO (GFP-CNO group) or saline (GFP-sal group), we observed a 14RS-dependent reduction in VGLUT2 expression to levels significantly lower than those of the unstressed GFP-CTR group (Figures 2B and 2C). In contrast, chemogenetic inhibition during RS (Gi-CNO group) prevented the VGLUT2 reduction observed after 14RS, whereas saline treatment (Gi-sal group) did not (Figures 2B and 2C). This intervention did not affect blood corticosterone levels (Figure S3A), confirming that stress was effectively induced despite chemogenetic inhibition. Further, we tested whether increased activity of VTAp–DCN neurons alone is sufficient to induce VGLUT2 reduction. Chemogenetic excitation alone (Gq-CNO group), achieved by expression of hM3Dq (Gq) in VTAp–DCN neurons and daily intraperitoneal CNO injections for 14 days in the absence of RS (Figure 2A), significantly reduced VGLUT2 levels in VTAp–DCN axons compared with the Gq-saline group (Figure 2D), whereas CNO administration in the absence of Gq expression had no effect (Figure S3B). Together, these results indicate that VGLUT2 reduction after chronic RS is activity-dependent in VTAp–DCN neurons.

**Figure 2.**
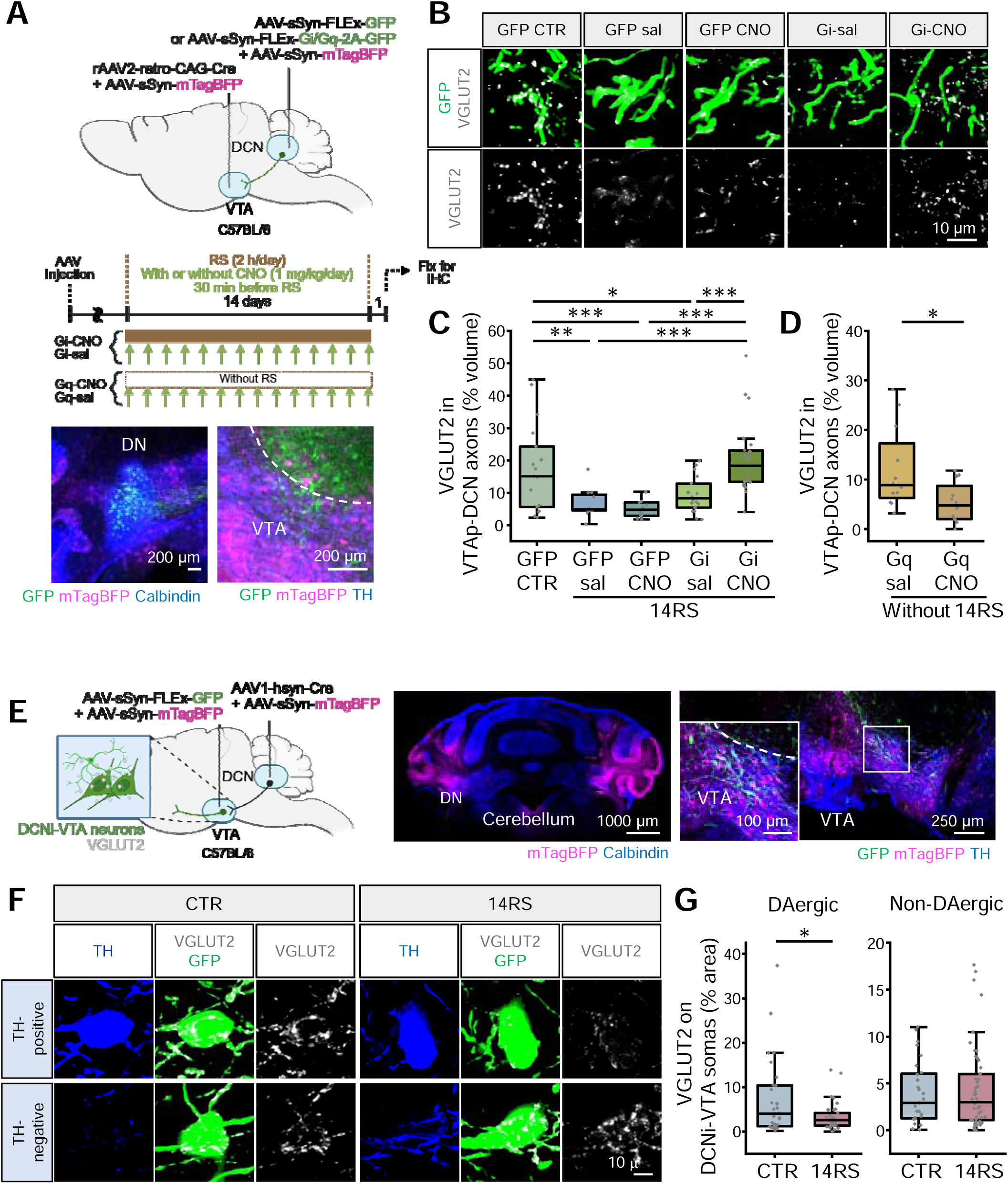
Activity-dependent VGLUT2 reduction in VTAp–DCN axons projecting to DAergic neurons. (A) Schematic of AAV injection and timeline for chemogenetic manipulation of VTAp–DCN neurons. CNO (1 mg/kg) was administered daily for 14 days with (Gi) or without (Gq) RS. Representative images of GFP and mTagBFP expression in the DCN (DN) and VTA are shown below. (B) Representative 3D projection images of GFP-labeled VTAp–DCN axons (green) and VGLUT2 staining (gray) restricted to GFP-positive VTAp–DCN axons in GFP-CTR, GFP-sal, GFP-CNO, Gi-sal, and Gi-CNO groups. (C) Percentage of VGLUT2-positive volume within VTAp–DCN axons in GFP-CTR (n = 17 slices from 8 mice), GFP-sal (n = 12 slices from 6 mice), GFP-CNO (n = 11 slices from 6 mice), Gi-sal (n = 18 slices from 8 mice), and Gi-CNO (n = 21 slices from 10 mice). (D) Percentage of VGLUT2-positive volume within VTAp–DCN axons in Gq-sal (n = 12 slices from 6 mice) and Gq-CNO (n = 12 slices from 6 mice). (E) Schematic of DCNi-VTA neuron labeling via AAV injection, along with representative images of mTagBFP expression in the cerebellum and GFP/mTagBFP expression in the VTA. The region outlined by the white square in the dorsolateral VTA is shown at higher magnification. (F) Representative 3D projection images of GFP-labeled DCNi-VTA neurons (green), TH (blue), and VGLUT2 (gray) restricted to GFP-positive regions in CTR and 14RS groups. (G) Percentage of VGLUT2-positive area in DCNi-VTA somata, quantified separately in TH-positive DAergic neurons (CTR: n = 34 somata from 10 mice; 14RS: n = 34 somata from 10 mice) and TH-negative non-DAergic neurons (CTR: n = 28 somata from 10 mice; 14RS: n = 50 somata from 10 mice). *p < 0.05, **p < 0.01, ***p < 0.001; one-way ANOVA followed by Fisher’s LSD post hoc test (C); two-tailed unpaired Student’s t-test (D and G).

Because the VTA comprises both DAergic and non-DAergic neurons, we next investigated which cell types are targeted by VTAp–DCN neurons exhibiting reduced VGLUT2 expression. Accordingly, we labeled VTA neurons receiving synaptic input from the DCN (DCNi-VTA neurons) with GFP (Figure 2E) by leveraging the anterograde transsynaptic spread of an AAV serotype 1 (AAV1) ^41^, and classified GFP-positive VTA neurons as DAergic or non-DAergic based on TH staining. Among all GFP-positive DCNi-VTA neurons, 47% were DAergic and 53% were non-DAergic (Figure S3C). Notably, a significant reduction in VGLUT2 was observed selectively in the vicinity of DAergic DCNi-VTA neurons (Figures 2F and 2G). These findings suggest that activity-dependent VGLUT2 reduction in VTAp–DCN neurons may preferentially affect DAergic neuron function in the dorsolateral VTA.

### Cerebellum-dependent changes in synaptic transmission in the VTA following chronic RS

Stress exposure can alter excitatory synaptic transmission in VTA DAergic neurons in a manner that depends on stress type and duration ^27,42–45^. Although VTAp–DCN neurons constitute only a subset of excitatory inputs to the dorsolateral VTA ^6,10^, we observed an activity-dependent reduction of VGLUT2 in these neurons specifically at synapses onto DAergic neurons. We therefore next examined whether chronic RS induces VTAp–DCN neuronal activity-dependent changes in excitatory synaptic transmission in DAergic VTA neurons. Miniature excitatory postsynaptic currents (mEPSCs) recorded from DAergic neurons in the dorsolateral VTA were compared between CTR and 14RS groups, with or without chemogenetic inhibition of VTAp–DCN neurons during RS (Figure 3A). DAergic neurons, identified either genetically in DAT-IRES-Cre;Ai6 mice (Figure 3B) or physiologically by detecting hyperpolarization-activated (Ih) currents (>15 pA in response to a voltage step from –60 mV to –110 mV; Figure S4A) as previously described ^46–52^, demonstrated a decrease in the frequency of mEPSCs in the 14RS group compared with the CTR group (Figures 3C, 3D, and S4B). This reduction was prevented, and mEPSC frequency even exceeded CTR levels, by chemogenetic inhibition of VTAp–DCN neurons during RS (Gi-CNO group; Figures 3C, 3D, and S4B). In contrast, the Gi-sal group exhibited a reduction in mEPSC frequency similar to that observed in the untreated 14RS group (Figures 3C, 3D and S4B). The amplitude of mEPSCs was comparable across all groups (Figures 3C, 3E, and S4C). These findings indicate that the reduction in mEPSC frequency in VTA DAergic neurons following chronic RS is mediated by RS-induced activation of VTAp–DCN neurons.

**Figure 3.**
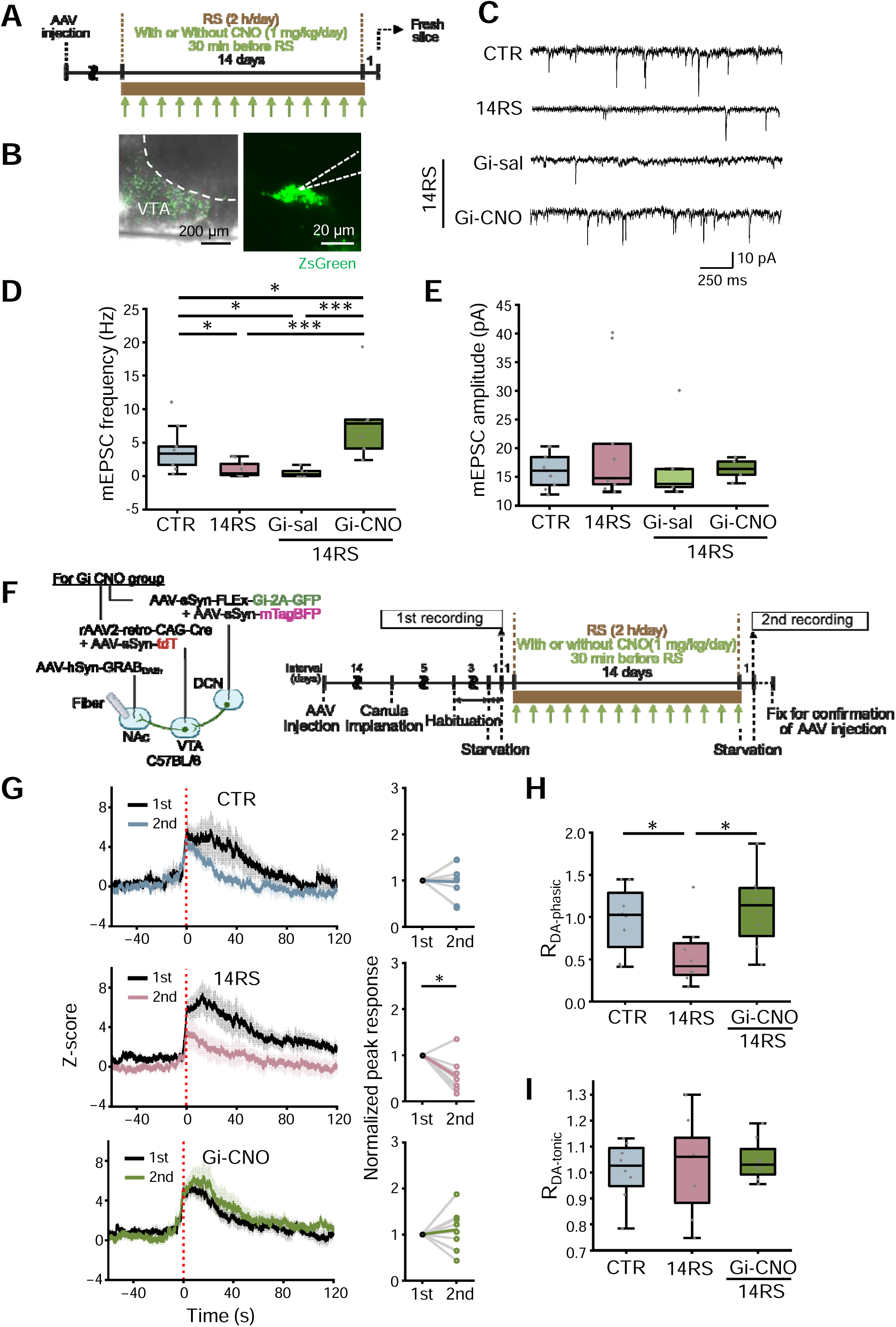
Dependence of 14RS-induced reductions in mEPSC frequency and DA release on VTAp–DCN neuronal activity. (A) Timeline of patch-clamp experiments following 14RS with or without chemogenetic inhibition. (B) ZsGreen expression in VTA DAergic neurons of a DAT-IRES-Cre;Ai6 mouse. (C) Representative mEPSC traces recorded from DAergic neurons in the CTR, 14RS, Gi-sal, and Gi-CNO groups. (D and E) Quantification of mEPSC frequency (D) and amplitude (E) in CTR (n = 9 cells from 5 mice), 14RS (n = 11 cells from 11 mice for (D); n = 10 cells from 10 mice for (E)), Gi-sal (n = 6 cells from 4 mice for (D); n = 5 cells from 3 mice for (E)), and Gi-CNO (n = 7 cells from 4 mice). Cells without detectable events were excluded from amplitude analysis. (F) Schematic of AAV injection and fiber implantation, and timeline for fiber photometry recording of DA release in the NAc. (G) Average z-scored GRAB_DA2h_ responses aligned to feeding onset (t = 0, red dotted line) in the first (black) and second (colored) recordings (left), and peak responses in the first and second recordings normalized to the first response for each mouse (right). For z-scored traces, solid lines indicate mean and shaded areas indicate SEM. (H) Second-to-first ratio of feeding-evoked peak DA responses (R_DA-phasic_). (I) Second-to-first ratio of baseline GRAB_DA2h_ signal (R_DA-Tonic_). n = 8 mice per group. *p < 0.05, ***p < 0.001; one-way ANOVA followed by Fisher’s LSD post hoc test (D, E, H and I); paired two-sample t-tests (G, right panels).

The seemingly paradoxical finding of a reduction in VGLUT2 but no decrease in mEPSC amplitude could be reconciled if there was also a reduction in the number of active synapses. Consistent with this idea, the number—but not the volume—of VGLUT2 puncta in VTAp–DCN axons was reduced after 14RS (Figure S4D). Thus, the decrease in mEPSC frequency may be attributable, at least in part, to a reduction in the number of active synapses, in line with reduced VGLUT2 expression.

### Cerebellum-dependent changes in DA release in the NAc after chronic RS

Chronic RS reduced VGLUT2 expression and excitatory synaptic transmission onto DAergic neurons in the dorsolateral VTA as a result of increased VTAp–DCN neuronal activity during RS. As dysregulation of the DA system has been widely implicated in depression ^20,25,53,54^, we hypothesized that DA release might also be altered after 14RS as a downstream consequence of VTAp–DCN neuronal activity. To identify brain regions targeted by DAergic DCNi-VTA neurons, we employed a dual AAV strategy in DAT-IRES-Cre mice (Figures S5A and S5B), injecting an AAV1 encoding Cre-dependent FlpO into the DCN and a FlpO-dependent GFP-expressing AAV into the VTA. Among several brain regions containing GFP-positive axons, robust GFP signals were observed in the nucleus accumbens (NAc), a major projection target of VTA DAergic neurons (Figure S5C). Consistent with a previous study ^55^, these results confirmed that DAergic DCNi-VTA neurons innervate the NAc.

Because GFP signals were more prominent in the shell of the NAc relative to the core, we performed fiber photometry primarily in the medial NAc shell (Figure S5D), following a previous study ^31^. DA release was measured using the genetically encoded DA sensor GRAB_DA2h_ ^56^ during feeding behavior ^31,57^ in CTR and 14RS mice (Figure 3F). Owing to inter-individual variability in GRAB_DA2h_ signals—likely stemming from differences in expression levels or fiber placement—DA levels were measured twice in each mouse to provide normalization, with or without RS applied during the 2-week interval between measurements (Figure 3F). Feeding elicited robust increases in DA signals in both groups (Figure 3G, top and middle). In the CTR group, feeding-evoked DA responses were comparable between the two time points, with no significant difference in peak values (Figure 3G, top). In contrast to the CTR group, 14RS mice exhibited a significant reduction in the peak value of the phasic DA response after 14RS compared with the pre-14RS measurement (Figure 3G, middle). Accordingly, the ratio of the second to the first response (R_DA-phasic_) was significantly lower in the 14RS group than in the CTR group (Figure 3H). Although comparisons of absolute GRAB_DA2h_ signal intensities across animals are unreliable, within-subject comparisons of baseline GRAB_DA2h_ signals can reflect changes in tonic DA levels. To assess this, we calculated the average GRAB_DA2h_ signal (normalized to the isosbestic signal) during the first 3 min of recording and derived the ratio of the second to the first measurement (R_DA-tonic_). R_DA-tonic_ did not differ significantly between the CTR and 14RS groups (Figure 3I), suggesting that chronic RS selectively reduced phasic, but not tonic, DA release in the NAc.

To determine whether the reduction in phasic DA release depended on VTAp–DCN neuronal activity, we chemogenetically inhibited these neurons during RS via AAV-mediated expression of Gi (Figure 3F). In the Gi-CNO group, feeding-evoked DA release remained unchanged even after 14RS (Figure 3G, bottom). Consistent with this, the R_DA-phasic_ in this group was comparable to that in CTR mice and significantly higher than that in untreated 14RS mice (Figure 3H). R_DA-tonic_ was similar across the CTR, 14RS, and Gi-CNO groups (Figure 3I). These findings indicate that the reduction in feeding-evoked phasic DA release in the NAc following chronic RS is mediated by VTAp–DCN neuronal activity.

### Contribution of VGLUT2 reduction to the emergence of depression-like behaviors

The temporal coincidence between VGLUT2 reduction and the emergence of depression-like behaviors, together with the dependence of both phenomena on VTAp–DCN neuronal activity, suggests a close association between VGLUT2 reduction and behavioral outcomes following chronic RS. To test this, we examined whether VGLUT2 overexpression (VGLUT2 OE) in VTAp–DCN neurons affects the emergence of depression-like behaviors. An AAV driving Cre-dependent expression of VGLUT2 and GFP via a 2A peptide was injected bilaterally into the DCN, and rAAV2-retro-CAG-Cre was injected into the VTA (Figure 4A), resulting in GFP-positive somata in the DCN and GFP-positive axons in the dorsolateral VTA (Figure S6A). GFP-positive axons were frequently associated with enlarged VGLUT2 puncta (Figure 4B), and puncta volume was significantly greater than that observed following injection of an AAV expressing GFP alone (Figures 4C and S6B), confirming that exogenously expressed VGLUT2 localized to axon terminals of VTAp–DCN neurons. Behavioral tests were conducted at the time points indicated in Figure S6C in four groups: GFP-CTR, GFP-14RS, VGLUT2 OE-CTR, and VGLUT2 OE-14RS. VGLUT2 OE had no detectable effect on behavior under unstressed CTR conditions, as GFP-CTR and VGLUT2 OE-CTR mice performed similarly across all behavioral assays (Figures 4D–4I). In contrast, GFP-14RS mice exhibited clear depression-like behaviors, including reduced grooming in the SST (Figure 4D), increased latency to feed in the NSFT (Figure 4E) without changes in home cage food consumption (Figure 4F), and increased immobility in both the TST (Figure 4G) and FST (Figure 4H), without alterations in general locomotor activity in the OFT (Figure 4I). Notably, these depression-like behaviors were significantly attenuated by VGLUT2 OE, as VGLUT2 OE-14RS mice performed comparably to CTR mice across all tests (Figures 4D–4I). These results indicate that VGLUT2 reduction in VTAp–DCN neurons contributes to the emergence of depression-like behaviors following chronic RS.

**Figure 4.**
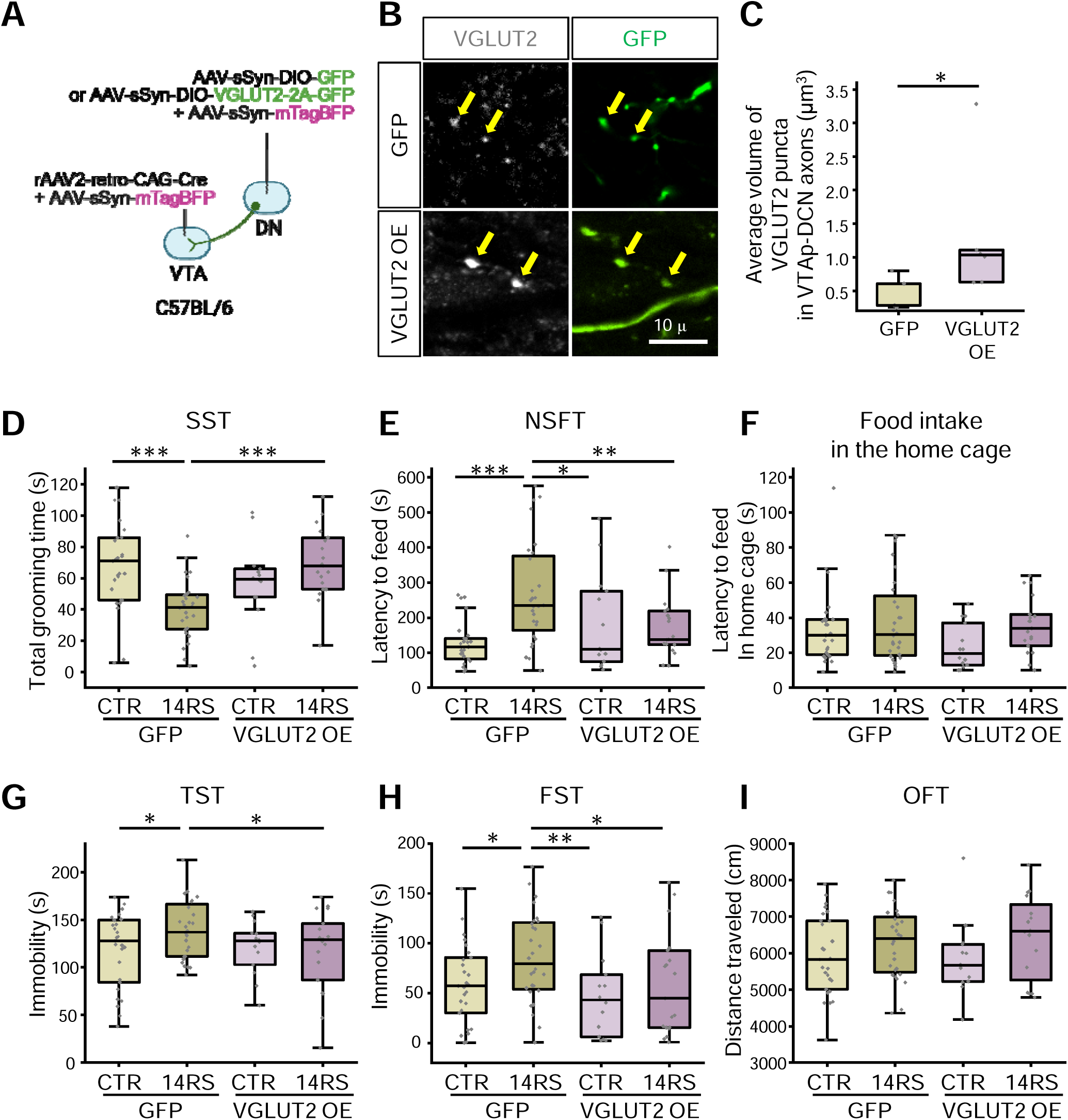
Attenuation of depression-like behaviors by VGLUT2 overexpression in VTAp–DCN neurons. (A) Schematic of AAV injection for selective VGLUT2 OE in VTAp–DCN axons. (B) Representative 3D projection images (10 μm thickness) of GFP (green) and VGLUT2 staining (gray). Yellow arrows indicate VGLUT2 puncta overlapping with GFP signals. (C) Average volume of VGLUT2 puncta within GFP-positive axons after GFP expression alone (n = 5 mice) or VGLUT2 OE (n = 6 mice). Data were obtained 2 weeks after AAV injection. (D–I) Total grooming time in the SST (D), latency to feed in the NSFT (E) and in the home cage (F), immobility time in the TST (G) and FST (H), and total distance traveled in the OFT (I), comparing GFP-CTR (n = 27 mice), GFP-14RS (n = 28 mice), VGLUT2 OE-CTR (n = 14 mice), and VGLUT2 OE-14RS (n = 19 mice in (D–H); n = 16 mice in (I)) groups. *p < 0.05, **p < 0.01, ***p < 0.001; Mann–Whitney U test (C); one-way ANOVA followed by Fisher’s LSD post hoc test (D–I).

### VGLUT2 reduction as an underlying mechanism for decreased synaptic transmission and DA release

We next examined whether the reductions in synaptic transmission in the VTA and DA release in the NAc following chronic RS occur downstream of VGLUT2 reduction in VTAp–DCN neurons. To this end, we performed electrophysiological recordings and fiber photometry experiments following the time course shown in Figure S6C in four groups: GFP-CTR, GFP-14RS, VGLUT2 OE-CTR, and VGLUT2 OE-14RS.

Under CTR conditions without RS, VGLUT2 OE alone had no effect on either the frequency or amplitude of mEPSCs recorded from DAergic neurons in the dorsolateral VTA, as these parameters were comparable between GFP-CTR and VGLUT2 OE-CTR groups (Figures 5A–5C). This suggests that VGLUT2 OE in VTAp–DCN neurons does not sufficiently increase synapse number or vesicle size under CTR conditions to be detected by mEPSC recordings, despite the increase in VGLUT2 puncta volume (Figure 4C). Nevertheless, VGLUT2 OE prevented the 14RS-induced reduction in mEPSC frequency, as no significant difference was detected between the VGLUT2 OE-14RS and CTR groups, although mEPSC frequency was not fully restored, as it also did not differ significantly from that of the GFP-14RS group (Figures 5A and 5B). In contrast, mEPSC amplitude remained comparable across all groups (Figures 5A and 5C). These results indicate that reduced excitatory synaptic transmission in dorsolateral VTA DAergic neurons following chronic RS is partially mediated by VGLUT2 reduction in VTAp–DCN neurons.

**Figure 5.**
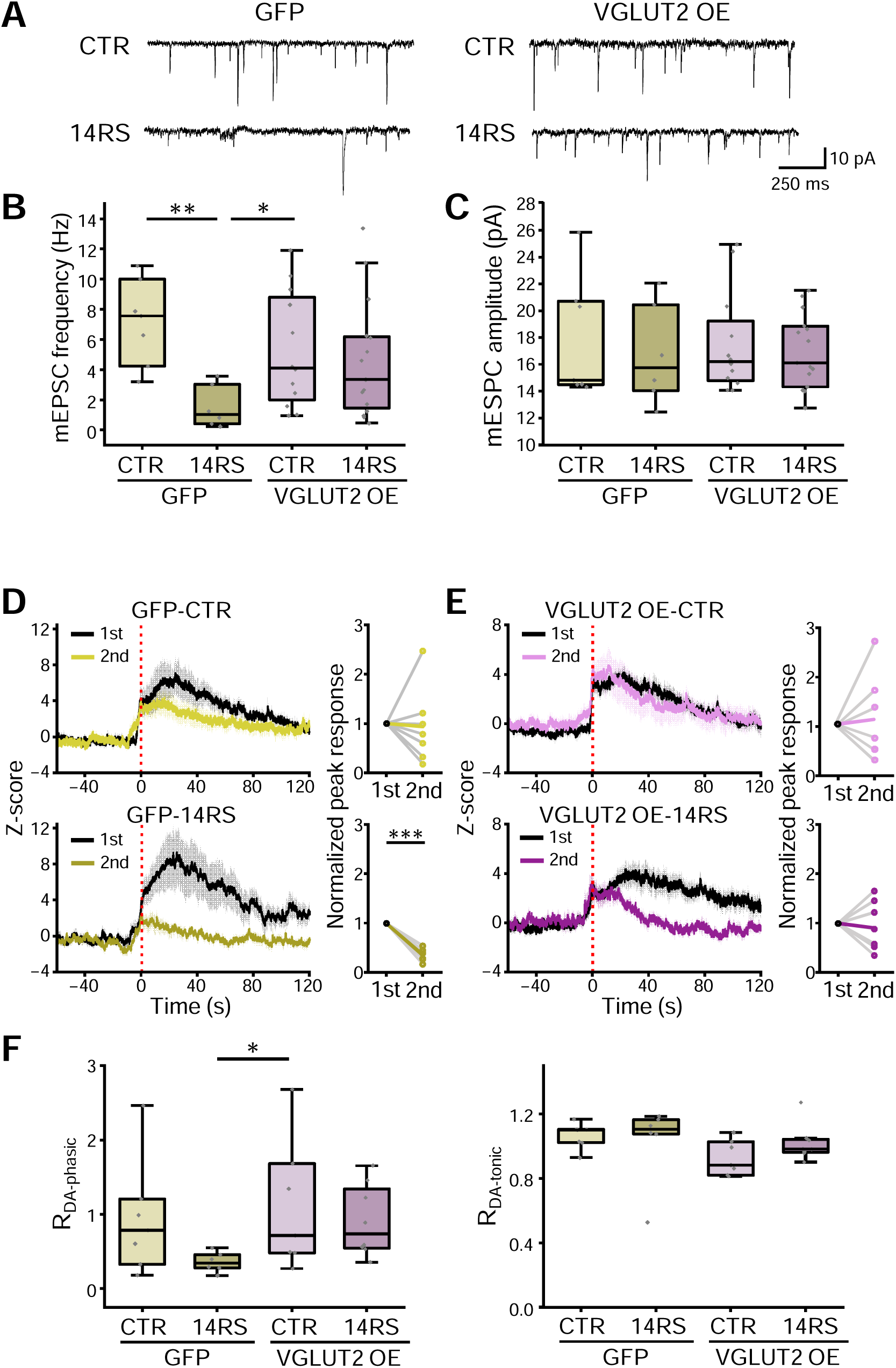
Prevention of 14RS-induced decreases in mEPSC frequency and DA release by VGLUT2 overexpression. (A) Representative mEPSC traces recorded from DAergic neurons in GFP-CTR, GFP-14RS, VGLUT2 OE-CTR, and VGLUT2 OE-14RS groups. (B and C) Quantification of mEPSC frequency (B) and amplitude (C) in GFP-CTR (n = 7 cells from 3 mice), GFP-14RS (n = 6 cells from 4 mice), VGLUT2 OE-CTR (n = 12 cells from 5 mice), and VGLUT2 OE-14RS (n = 16 cells from 8 mice for (B); n = 15 cells from 8 mice for (C)). Cells without detectable events were excluded from amplitude analysis. (D and E) Average z-scored GRAB_DA2h_ responses aligned to feeding onset (t = 0, red dotted line) in the first (black) and second (colored) recordings (left), and peak responses normalized to the first response for each mouse (right). For z-scored traces, solid lines indicate mean and shaded areas indicate SEM. (F) Second-to-first ratio of feeding-evoked peak DA responses (R_DA-phasic_, left) and second-to-first ratio of baseline GRAB_DA2h_ signal (R_DA-Tonic_, right). For fiber photometry recordings (D–F), n = 7, 6, 7, and 8 mice for GFP-CTR, GFP-14RS, VGLUT2 OE-CTR, and VGLUT2 OE-14RS groups, respectively. *p < 0.05, **p < 0.01, ***p < 0.001; one-way ANOVA followed by Fisher’s LSD post hoc test (B, C and F); paired two-sample t-tests (D and E, right panels).

Fiber photometry measurements of DA release in the NAc showed that peak values of feeding-evoked DA release in the GFP-CTR group were comparable between the two time points separated by a 2-week interval without RS, whereas peak DA release in the GFP-14RS group was significantly reduced after 14RS compared with the pre-14RS measurement (Figure 5D). In contrast, peak DA release in both the VGLUT2 OE-CTR and VGLUT2 OE-14RS groups remained unchanged between the two time points (Figure 5E), indicating that VGLUT2 OE prevented the 14RS-induced decrease in DA release. Consistent with this, R_DA-phasic_ in the VGLUT2 OE-14RS group was comparable to that in the GFP-CTR and VGLUT2 OE-CTR groups (Figure 5F). R_DA-phasic_ in the GFP-14RS group was reduced relative to the other groups, although statistical significance was reached only in comparison with the VGLUT2 OE-CTR group (Figure 5F). R_DA-tonic_ was similar across all groups, indicating that tonic DA release was not affected by either 14RS or VGLUT2 OE. Together, these results support the conclusion that the decrease in NAc DA release following chronic RS is mediated, at least in part, by VGLUT2 reduction in VTAp–DCN neurons.

## Discussion

While we have previously demonstrated that VTAp–DCN neurons proactively regulate the development of depression-like behaviors ^10^, here we elucidate the underlying mechanisms, as summarized in Figure 6. We found that VGLUT2 expression was reduced in the axons of VTAp–DCN neurons as a result of chronic stress, with a temporal profile that coincided with the emergence of depression-like behaviors. This reduction was preferentially observed in axons projecting to DAergic neurons. Consistent with this decrease, mEPSC frequency in dorsolateral VTA DAergic neurons, as well as DA release in the NAc, was reduced following chronic RS. Importantly, these reductions in VGLUT2 expression, excitatory synaptic transmission, and DA release were dependent on the activity of VTAp–DCN neurons. Notably, preventing the RS-induced reduction of VGLUT2 through targeted overexpression in VTAp–DCN neurons protected stressed mice from exhibiting depression-like behaviors and was accompanied by recovery of excitatory synaptic transmission and NAc DA release. Together, these results indicate that reduced VGLUT2 levels in VTAp–DCN neurons contribute to chronic RS-induced depression-like behaviors. In light of a previous report showing that optogenetic inhibition of VTA DAergic neurons acutely induces depression-like behaviors ^23^, impaired DA release may directly contribute to these behaviors. Thus, the reductions in VGLUT2 expression, excitatory synaptic transmission, and DA release observed in the present study likely represent a sequence of events underlying the emergence of depression-like behaviors.

**Figure 6.**
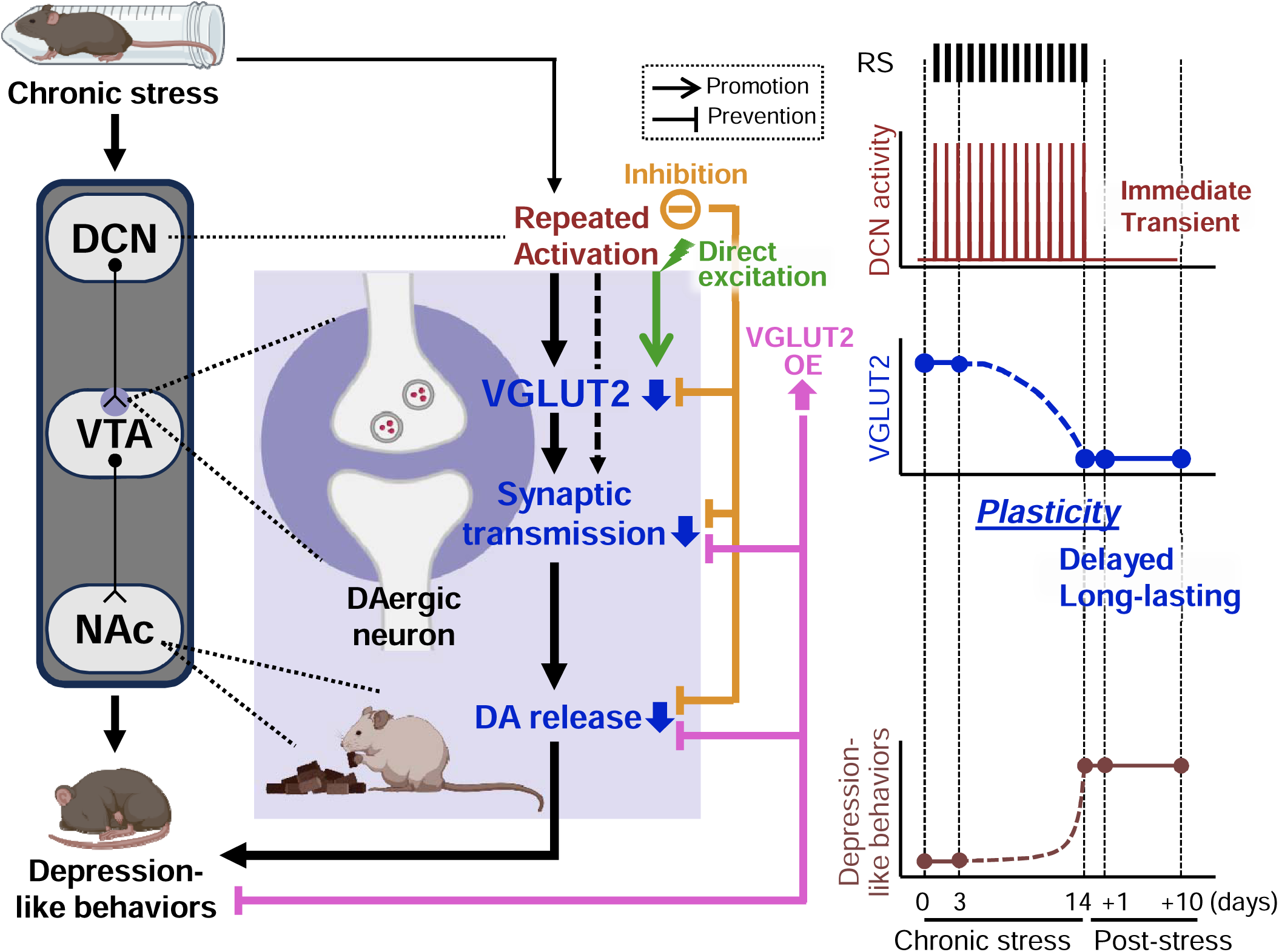
Diagram summarizing the results of this study. We showed that chronic RS reduced VGLUT2 in VTAp–DCN neurons, temporally coinciding with the emergence of depression-like behaviors. Along with VGLUT2 reduction, excitatory synaptic transmission onto VTA DAergic neurons and DA release in the NAc were also decreased, and manipulation of VTAp–DCN neurons further supported a causal relationship among these changes. Based on these findings, we propose that chronic stress–dependent repeated activation of VTAp–DCN neurons induces a delayed and persistent reduction of VGLUT2 in their axons, leading to depression-like behaviors, likely through reduced synaptic transmission onto VTA DAergic neurons and diminished DA release in the NAc.

### Temporal aspects of RS-triggered events

Numerous studies have shown that 14RS induces depression-like behaviors ^10,32–35^. In these studies, such behaviors persisted for one to two weeks after the cessation of RS, consistent with the long-lasting symptoms observed in patients with major depressive disorder. In contrast, acute or short-duration RS has been reported to induce other behavioral changes, such as anxiety-like behaviors or memory impairments ^58–62^, and is generally considered insufficient to induce depression-like behaviors. Consistent with this view, we did not observe clear depression-like behaviors following 3RS.

RS triggers a range of biological changes with distinct temporal dynamics, which may differentially influence behavioral outcomes. For example, blood corticosterone levels rise rapidly following stress exposure and mediate stress responses through complex mechanisms ^63^, contributing to both acute stress coping and longer-term maladaptive effects associated with chronic elevation ^64,65^. Because DCN neuronal activity also increases rapidly upon RS exposure—similar to corticosterone elevation—it is possible that DCN activity contributes not only to chronic RS-induced depression-like behaviors ^10^, but also to behavioral or biological responses associated with acute RS. Given that DAergic regulation plays a central role in stress responses ^25,66^ and that direct DN stimulation can evoke DA release in the NAc ^30,31^, VTAp–DCN neurons may modulate DAergic activity during acute RS and thereby participate in stress-coping processes.

The temporal discrepancy between rapid DCN activation and the delayed onset of depression-like behaviors supports the idea that VTAp–DCN neurons contribute to these behaviors through repeated activation that induces slow, persistent plasticity in downstream circuits. Among the synaptic molecules examined, VGLUT2 reduction in VTAp–DCN axons exhibited a temporal profile consistent with this model. In addition, experiments using chemogenetic inhibition and excitation demonstrated that this VGLUT2 reduction depends on repeated activation of VTAp–DCN neurons, supporting VGLUT2 reduction as a plastic component of this model. Moreover, preventing VGLUT2 reduction through targeted overexpression suppressed the emergence of depression-like behaviors following chronic RS, indicating the its involvement in these behaviors. Together, these converging findings—temporal coincidence, activity dependence, and causal relevance—establish VGLUT2 reduction as a key plastic alteration linking rapid VTAp–DCN neuronal activity to delayed depression-like behaviors, potentially contributing directly to their manifestation.

### Mechanisms linking VGLUT2 reduction to depression-like behaviors

While our data show a clear association between reduced VGLUT2 expression in VTAp–DCN neurons and depression-like behaviors, the mechanisms linking these changes are not immediately apparent. Disruption of DAergic regulation is broadly implicated in depression ^20,25,53,54^. Notably, bidirectional optogenetic manipulation of VTA DAergic neurons has demonstrated a tight relationship between reduced DAergic control in the NAc and depression-like behaviors ^23^. Here we found that chronic RS-induced impairment of NAc DA release was driven, at least in part, by activity-dependent VGLUT2 reduction in VTAp–DCN neurons. Thus, reduced DA release likely represents a critical link between VGLUT2 reduction in VTAp–DCN neurons and depression-like behaviors. At the single-cell level, we also observed that chronic RS decreased excitatory synaptic transmission in dorsolateral VTA DAergic neurons, partially dependent on the VGLUT2 reduction. This weakening of synaptic transmission likely underlies impaired DA release in the NAc, given the importance of afferent modulation of the DAergic system in depression-like behaviors^54^. Together, these findings support a model in which chronic RS leads to activity-dependent VGLUT2 reduction in VTAp–DCN neurons, resulting in weakened excitatory synaptic input onto DAergic neurons, impaired DA release, and ultimately depression-like behaviors.

### Plasticity regulated by cerebellar efferent pathways

The cerebellum is classically recognized for its role in coordinating the timing of motor actions. Consistent with this role, DCN output neurons exhibit activity patterns that are tightly aligned with motor execution ^67–69^. These temporally precise activity patterns can be modified by associative learning paradigms that pair conditioned and unconditioned stimuli, suggesting that DCN activity reflects plastic changes in the cerebellar cortex and contributes directly to learned motor behaviors. Beyond motor control, DCN neuronal activity also contributes to the expression of learned cognitive behaviors. For example, selective silencing of DCN neurons during testing impaired performance in a forced alternation task even after the learning phase was completed ^5^. In addition, specific DCN projections have been shown to regulate emotional behaviors during conditioning, including cued fear memory formation and the anxiolytic effects of rotarod training ^8,11,13,70^. These findings suggest that DCN outputs induce plastic changes in target regions during conditioning, thereby shaping learned emotional states. Indeed, conditioning stimuli such as cued fear or rotarod training have been shown to elicit synaptic plasticity in brain regions innervated by DCN neurons ^11,70^. In the present study, chronic RS induced plastic alterations at VTA synapses—specifically, reductions in VGLUT2 expression and excitatory synaptic transmission—dependent on DCN neuronal activity. Taken together with previous findings, these results suggest that cerebellar outputs regulate behavior across a broad temporal range, from influencing real-time responses through precisely timed signaling to contributing to delayed and persistent behavioral changes by conveying contextual information and inducing plasticity in downstream circuits.

Accumulating evidence links cerebellar abnormalities to stress-related psychiatric disorders, including major depressive disorder and post-traumatic stress disorder (see reviews ^71,72^). Building on our previous finding that VTAp–DCN neurons proactively regulate depression-like behaviors ^10^, the present study identifies downstream mechanisms that give rise to delayed and persistent behavioral outcomes. An open question concerns the nature of the information conveyed by VTAp–DCN neurons. Although these neurons are activated during RS—suggesting the transmission of aversive signals—previous studies have also associated VTAp–DCN neuronal activity with reward and satiety ^6,31^. This duality is consistent with the heterogeneous organization of VTA DAergic neurons, which respond to both rewarding and aversive stimuli ^22,24,25^. VTAp–DCN neurons may therefore represent an upstream pathway contributing to this functional diversity. Understanding the general role of VTAp–DCN neurons within the broader regulatory network, as well as the specific information they transmit during the development of depression-like behaviors, will be an important goal for future studies. The latter may also provide crucial insight into how VTAp–DCN neurons, together with the downstream mechanisms identified in this study, cooperate with other neuronal circuits involved in chronic RS-induced depression-like behaviors. Regardless of the specific information conveyed, our present study demonstrates that repeated activation of VTAp–DCN neurons induces plastic changes in the VTA, leading to reduced DA release, which is likely directly associated with depression-like behaviors such as anhedonia and behavioral despair.

### Limitations of this study

We propose that reduced VGLUT2 expression in VTAp–DCN neurons is associated with the manifestation of depression-like behaviors. The temporal coincidence between VGLUT2 reduction and behavioral outcomes supports this proposal. However, because we did not control the timing of VGLUT2 overexpression, the phase specificity of this reduction—whether it contributes to the manifestation phase, the development phase, or both—remains unresolved. This issue could be addressed by temporally controlling VGLUT2 overexpression using a tetracycline-controlled system, although such a system may lack sufficient temporal precision. Ideally, future studies should develop tools enabling more precise temporal manipulation of axonal VGLUT2 expression—such as optogenetic approaches including LARIAT ^73^ or CALI ^74^—to test whether manipulation during specific phases differentially affects the development or expression of depression-like behaviors.

In the present study, chronic RS-induced decreases in excitatory synaptic transmission were assessed using mEPSC recordings, a technically straightforward approach that enables data acquisition from a large number of mice across multiple experimental conditions (CTR, 14RS, Gi-sal, Gi-CNO, GFP-CTR, GFP-14RS, VGLUT2 OE-CTR, and VGLUT2 OE-14RS). Nevertheless, the mechanisms by which reduced VGLUT2 expression leads to decreased mEPSC frequency require careful consideration. Reduced mEPSC frequency is likely attributable, at least in part, to a decrease in the number of functional synapses resulting from VGLUT2 downregulation, consistent with our observation of a reduced number of VGLUT2 puncta. In addition, given reports that VGLUT expression levels correlate with release probability ^75^, reduced release probability may also contribute. Importantly, previous studies have shown that optogenetic stimulation of DCN neurons evokes EPSCs in only a subset of VTA neurons—approximately 25% ^10^ or 50% ^6^. Although these estimates may be conservative, they suggest that only a fraction of the excitatory inputs to DAergic neurons in the dorsolateral VTA originates from VTAp–DCN neurons. Thus, although the observed decrease in mEPSC frequency likely reflects weakened DCN-derived transmission via VGLUT2 downregulation, contributions from other glutamatergic inputs cannot be excluded. Even so, these inputs are likely indirectly regulated by VTAp–DCN neurons, as inhibition of these neurons fully attenuated the RS-induced decrease in mEPSC frequency. One possible mechanism is that activity of VTAp–DCN neurons engages endocannabinoid-mediated retrograde signaling; such signaling has been reported to regulate excitatory inputs onto VTA DAergic neurons ^76–78^. Elucidating the contribution of additional glutamatergic inputs, the underlying mechanisms, and their behavioral relevance will require further methodological and conceptual advances.

## Supporting information

Table S1. Statistics

## Acknowledgments

We thank members of the Tanaka-Yamamoto and Yamamoto laboratories for valuable discussions during the project, and Dr. Yulong Li for kindly providing us with the plasmids. Schematic figures were created using BioRender.com. This work was supported by the Korea Institute of Science and Technology Institutional Program (Project no.: 26Z9001), the National Research Foundation of Korea (NRF) grant funded by the Korean Ministry of Science and ICT (MSIT) (NRF grant nos.: RS-2021-NR059790, RS-2022-NR070223, and RS-2024-00338607).

## Author contributions

Conceptualization, Y.Y. and K.T.-Y.; Methodology, S.K., T.K., D.K. S.-J.B., T.J.M., Y.Y. and K.T.-Y.; Investigation, S.K., T.K., and D.K.; Funding acquisition, Y.Y. and K.T.-Y.; Resources, T.J.M.; Supervision, Y.Y. and K.T.-Y.; Writing—original draft, S.K. and K.T.-Y.; Writing—review and editing, S.K., T.K., D.K., S.-J.B., T.J.M., Y.Y. and K.T.-Y.

## Competing interests

The authors declare that they have no competing interests.

## Data and materials availability

All data needed to evaluate the conclusions in the paper are present in the paper and/or the Supplemental Information.

## Supplemental information

Figures S1–S6 and Table S1

## Materials and Methods

### Mice

All experiments involving mice were performed in accordance with the Institutional Animal Care and Use Committee of Korea Institute of Science and Technology. C57BL/6J mice were used for behavioral tests, immunohistochemistry, and fiber photometry. DAT-IRES-Cre (JAX #006660, B6.SJL-Slc6a3tm1.1(cre)Bkmn/J) mice were used for neuronal tracing analysis. Electrophysiological experiments were conducted using C57BL/6J and DAT-IRES-Cre;Ai6 mice, and western blot analysis was performed using DAT-IRES-Cre;Ai6 mice. DAT-IRES-Cre and Ai6 ZsGreen reporter mice (B6.Cg-Gt(ROSA)26Sortm6(CAG-ZsGreen1)Hze/J) were crossed to obtain DAT-IRES-Cre;Ai6 mice. Since our previous study ^10^ showed that chronic RS-induced depression-like behaviors were similar in both sexes, male C57BL/6J mice were used in the present study. To maximize experimental efficiency, DAT-IRES-Cre;Ai6 mice of both sexes were used, with same-sex littermates randomly assigned to different experimental groups. All mice were group-housed in same-sex groups of up to 6 mice per cage, under a 12-h light/dark cycle, with ad libitum access to food and water and a controlled temperature of 23–25°C. The ages of the mice are described in the individual sections explaining the different procedures.

### AAV production and stereotaxic injection

AAV serotype 1 (AAV1) and rAAV2-retro were used for anterograde and retrograde labeling, respectively. AAV1 was also used for transsynaptic anterograde tracing. AAV constructs were generated by cloning using the following plasmids: pAAV-sSyn and pAAV-sSyn-FLEx from our previous study ^79^; pAAV-hSyn derived from pAAV-hSyn-DIO-hM3D(Gq)-mCherry (Addgene, #44361); and pAAV-sSyn-DIO and pAAV-sSyn-FLEx(FRT), constructed using a DIO cassette from the same Addgene plasmid and a FLExFRT cassette from pAAV-hSyn-FLExFRT-mGFP-2A-Synaptophysin-mRuby (Addgene, #71761). The cDNA fragments for FlpO, VGLUT2 (GenBank accession number: BC038375), and NLS-Cre-Myc were obtained by PCR from pAAV phSyn1(S)-FlpO-bGHpA (Addgene, #51669), cDNA synthesized from mouse brain RNA, and pCAG-Cre (Addgene, #13775), respectively. The pAAV-hSyn-GRAB_DA2h plasmid (Addgene, #140554, ^56^) was kindly provided by Dr. Yulong Li (Peking University). Additional plasmids used in our previous studies—including pAAV-sSyn-FLEx-hM4Di(Gi)-2A-GFP, pAAV-sSyn-FLEx-hM3Dq(Gq)-2A-GFP, pAAV-sSyn-tdT, pAAV-sSyn-mTagBFP, paavCAG-iCre (Addgene, #51904), and pAAV-sSyn-FLEx-GFP ^10^—were also employed. AAV vectors with estimated titers from 10^12^–10^13^ vector genome copies were produced as described previously ^79^.

For stereotaxic injections, mice were anesthetized with Avertin (250 μg/g body weight) and injected using a stereotaxic apparatus (Narishige or Kopf) together with a microinjection pump (Nanoliter 2010, WPI, Inc). The volume of AAV injected was 0.5 μL in the DCN, 0.5 μL in the VTA, and 0.7 μL in the NAc. Injections into the DCN were primarily targeted to the DN, as VTAp–DCN neurons predominantly originate from the DN ^10^. Stereotaxic coordinates were as follows: DCN, anteroposterior (AP) −6.00 mm, mediolateral (ML) ±2.30, ventral (V) −2.60 mm relative to bregma; VTA, AP −3.1 mm, ML 0 mm, V −4.20; NAc, AP 0.98 mm, ML ±0.75, and V −4.20 mm. Mice subjected to the AAV injections were 3–4 weeks old for electrophysiological analysis and 5–6 weeks old for other analyses. The specific AAV vectors used for each analysis are depicted in the schematic diagrams of the figures. In brief, DCN axons in the VTA were visualized by unilateral injection of AAV-sSyn-tdT into the DCN. To achieve targeted expression of Gi, Gq, GFP, or VGLUT2 in VTAp–DCN neurons, rAAV2-retro-CAG-iCre was injected into the VTA, and a Cre-dependent AAV encoding Gi (AAV-sSyn-FLEx-Gi-GFP), Gq (AAV-sSyn-FLEx-Gq-GFP), GFP (AAV-sSyn-FLEx-GFP or AAV-sSyn-DIO-GFP), or VGLUT2 (AAV- sSyn-DIO-VGLUT2-2A-GFP) was injected into the DCN. To label DCNi-VTA neurons, AAV1-hSyn-Cre and AAV-sSyn-FLEx-GFP were injected into the DCN and VTA of C57BL/6J mice, respectively. For these analyses using VTAp–DCN neuron manipulation or DCNi-VTA neuron labeling, bidirectional DCN injections were combined with a midline VTA injection. To specifically label DAergic DCNi-VTA neurons, unilateral injection of AAV-sSyn-FLEx-FlpO into the DCN was combined with injection of AAV-sSyn-FLEx(FRT)-GFP into the VTA of DAT-IRES-Cre mice. For fiber photometry measurement of DA release in the NAc, AAV-hSyn-GRAB_DA2h_ was unilaterally injected into the NAc. AAV-sSyn-mTagBFP was occasionally used as an injection marker. After the surgery, mice were kept on a heating pad until recovery from anesthesia and then returned to their home cages.

### Restraint stress

RS was delivered as described previously ^10^. In brief, 5–6-week-old or 8-week-old mice were placed individually in a well-ventilated 50 mL conical tube for electrophysiological or other analyses, respectively. To minimize excess space, an 8.5–9.5 cm tube insert was placed inside the 50 mL tube after the mouse, with the size adjusted for each animal. Mice were subjected to 2 h of RS daily for 3 (3RS) or 14 (14RS) consecutive days. CTR mice not exposed to RS were age-matched to those in the 14RS group. For chemogenetic inhibition during RS, CNO stock solution (100 mM in dimethyl sulfoxide) was diluted in saline and administered intraperitoneally (100 μL/mouse) at 1 mg/kg. Control mice received saline only. CNO or saline was administered 30 min before RS. To examine the effects of chemogenetic excitation alone, mice expressing Gq received daily intraperitoneal injections of CNO or saline for 14 consecutive days without RS.

Throughout this study, “3RS” and “14RS” in the figures refer to results obtained from samples collected and experiments conducted or initiated one day after the final RS session, whereas “14RS+10” refers to results obtained 10 days after the final RS session, unless otherwise specified.

### Corticosterone ELISA

Mice were anesthetized with isoflurane, and blood samples were collected by cardiac puncture 30 min after the end of 3RS (3RS Imm) or 14RS (14RS Imm), at 8.5 or 10 weeks of age, respectively. Blood samples were kept at room temperature for 30 min and centrifuged at 2200 rpm for 20 min to obtain serum, which was stored at −20 °C until use. Serum samples were assayed using a corticosterone ELISA kit (ADI-901-097, Enzo Life Sciences) according to the manufacturer’s instructions. Corticosterone levels were quantified using a microplate reader (M200PRO, TECAN).

### Behavioral analysis

To evaluate depression-like behaviors, four behavioral tests—SST, NSFT, TST, and FST—were conducted. The OFT was used to assess general locomotor activity. All behavioral tests were performed in a sound-attenuated room between 10:00 and 18:00. Mice were habituated to the testing room for 30 min before each test. Before behavioral testing, mice were handled for 5 min per day for 3 consecutive days during the final 3 days of RS. Behavioral testing began with the OFT one day after completion of 3RS or 14RS, at 8.5 or 10 weeks of age, respectively, followed by the SST and NSFT on the next two consecutive days. The TST and FST were performed one week later. Thus, behavioral testing was completed approximately 10 days after the end of RS. Behaviors were video-recorded and analyzed post hoc using EthoVision software (Noldus).

The number of mice used for different behavioral analyses sometimes varied due to unexpected mouse deaths and recording errors in the OFT. An additional set of TST and OFT experiments using the CTR and 14RS groups was conducted for experimental conditioning, and these data were also included. The exact numbers of mice used are described in the figure legends and in Table S1.

#### Open Field Test (OFT)

The OFT was conducted to confirm that increased immobility observed in the TST and FST was not due to reduced locomotion. Each mouse was placed in the center of an arena (30 × 30 × 30 cm) under 20 lux illumination and allowed to explore for 30 min. Movements were recorded by an overhead camera, and total distance traveled was analyzed.

#### Sucrose Splash Test (SST)

Each mouse was placed in its home cage with familiar bedding and sprayed with 200 μL of 10% sucrose solution on its dorsal coat. Grooming behavior was video-recorded, and total grooming duration was measured. Grooming behaviors included facial washing with the forepaws, body licking, and fur scrubbing to remove the sucrose solution. Very brief grooming episodes (< 1 s) were excluded from analysis.

#### Novelty Suppressed Feeding Test (NSFT)

After 24 h of food deprivation in the home cage, each mouse was placed in a square open-field arena (50 × 50 × 35 cm) with new bedding. A single food pellet soaked in 50% sucrose was placed in the center under bright illumination (800 lux). Latency to feed was measured for up to 10 min. Mice that did not eat within 10 min were excluded from analysis. After the test in open-field arena, mice were immediately returned to their home cages, and latency to feed was recorded for an additional 5 min to verify feeding motivation.

#### Tail Suspension Test (TST)

Each mouse was suspended by attaching the tip of its tail to a horizontal rod 50 cm above the floor using adhesive tape. To prevent climbing, a 6-cm segment of a 15 mL conical tube was placed around the tail. Behavior was recorded for 6 min, and the last 4 min were analyzed for immobility using EthoVision software with an immobility threshold of 3%. Immobility was defined as complete absence of movement while suspended.

#### Forced Swimming Test (FST)

Each mouse was placed in an acrylic cylinder (30 cm high, 15 cm diameter) filled with water (22–24°C) to a depth of 15 cm. Behavior was recorded for 6 min, and the last 4 min were analyzed for immobility using EthoVision software with an immobility threshold of 3%. Immobility was defined as remaining motionless except for movements necessary to float and maintain balance or keep the head above water.

### Western blot, immunohistochemistry, and confocal imaging

Primary antibodies used were rabbit anti-GluR1 (ab31232, Abcam), mouse anti-GluR2 (MAB397, Millipore), rabbit anti-GluR2 phosphorylated at Ser880 (p-GluR2) (ab52180, Abcam), guinea pig anti-VGLUT2 (AB2251-I, Millipore), rabbit anti-PSD-95 (ab18258, Abcam), rabbit or mouse anti-TH (AB152 and MAB318, Millipore), rabbit anti-calbindin (C2724, Sigma-Aldrich), rabbit anti-c-Fos (2250, Cell Signaling Technology), and mouse anti-β-actin (sc-47778, Santa Cruz). Secondary antibodies included HRP-conjugated anti-mouse or anti-rabbit IgG (GE Healthcare or Invitrogen), HRP-conjugated anti-guinea pig IgG (Jackson ImmunoResearch Laboratories), and Alexa Fluor 488- or 647-conjugated anti-mouse, anti-rabbit, or anti-guinea pig IgG antibodies (Thermo Fisher Scientific).

For both western blot and immunohistochemical analyses of synaptic molecules, samples were collected one day after 3RS or 14RS, or 10 days after 14RS, at 8.5, 10, or 11.5 weeks of age, respectively. For confirmation of VGLUT2 OE, samples were collected 2 weeks after injection of AAV-sSyn-DIO-GFP or AAV-sSyn-DIO-VGLUT2-2A-GFP, at 8 weeks of age.

For western blot analysis, DAT-IRES-Cre;Ai6 mice were deeply anesthetized with isoflurane and decapitated. Coronal midbrain slices (300 μm) containing the VTA were prepared using a vibrating microtome (Leica VT1200S) in ice-cold normal artificial cerebrospinal fluid (aCSF) containing (in mM): 125 NaCl, 25 NaHCO_3_, 2.5 KCl, 1.25 NaH_2_PO_4_, 11 glucose, 1.3 MgCl_2_, and 2.5 CaCl_2_. ZsGreen-positive VTA regions were visualized using a fluorescence flashlight and filter glasses and were punched using a 0.5 mm puncher (Palkovits brain punch, Stoelting) (see Figure S1A). Two punches from the left and right VTA of one mouse (approximately 25 μg in total) were transferred into 50 μL SDS-PAGE sample buffer. Because VTA samples were collected sequentially from slices, the number of mice that could be processed in one experimental set was limited. Therefore, in each experimental set, one RS group (3RS, 14RS, or 14RS+10) was compared with the CTR group, with 3–5 mice per group. After centrifugation at 5,000 rpm for 3 min, supernatants were directly subjected to immunoblotting to detect VGLUT2, GluR2, and p-GluR2 to prevent heat-induced signal reduction. The remaining supernatants were boiled at 95°C for 5 min and used to detect GluR1, PSD-95, TH, and β-actin. Band intensities and background signals were quantified using Fiji (an open-source distribution of ImageJ; ^80^). After background subtraction, signals for GluR1, GluR2, p-GluR2, VGLUT2, PSD-95, and TH were normalized to the corresponding β-actin signals.

For immunohistochemistry, mice were anesthetized with isoflurane and perfused transcardially with 4% paraformaldehyde (PFA) in 0.1 M sodium phosphate buffer (pH 7.4). Brains were immediately removed and post-fixed in 2% PFA overnight at 4°C. Coronal slices of the cerebellum (100 μm) and midbrain including the VTA (50 μm) were prepared using a vibratome (Leica VT1200S). Slices were blocked for 30 min at room temperature in 5% normal goat serum in PBS, incubated with primary antibodies overnight at 4°C, washed, and incubated with secondary antibodies overnight at 4°C. Slices were mounted and imaged using an A1R or AX laser-scanning confocal microscope (Nikon) with 10× (1,024 × 1,024 resolution) and 60× (1,024 × 1,024 resolution) objectives.

To identify the primary projection targets of DAergic DCNi-VTA neurons, these neurons were labeled with GFP by injecting AAV1-sSyn-FLEx-FlpO into the DCN and AAV-sSyn-FLEx(FRT)-GFP into the VTA of 5-week-old DAT-IRES-Cre mice. Five to six weeks later, mice were perfused as described above. To examine GFP-positive axons, every other 100 μm coronal section throughout the whole brain was prepared using a vibratome (Leica VT1200S). Low-magnification images were collected from known projection targets of VTA DAergic neurons, including the NAc, thalamus, and hippocampus.

In mice used for fiber photometry and behavioral experiments following AAV injection, expression of GFP, mTagBFP, tdT, or GRAB_DA2h_, and fiber placement were confirmed after completion of the experiments.

### Image analyses

Image analyses of GluR2 and VGLUT2 were performed after acquiring single-field-of-view z-stack images (212 × 212 × 25 μm or 295 × 295 × 25 μm; 50 slices at 0.5 μm intervals) from the dorsolateral VTA using NIS-Elements (Nikon) and Fiji software. One to three slices per mouse were used for analysis. The dorsolateral VTA was identified based on TH immunostaining and fluorescent labeling after AAV injection. When AAV-sSyn-tdT, AAV-sSyn-FLEx/DIO-GFP, AAV-sSyn-FLEx-Gi/Gq-2A-GFP, or AAV-sSyn-DIO-VGLUT2-2A-GFP was injected into the DCN, regions containing both TH-positive DAergic neurons and tdT- or GFP-positive axons were analyzed. When DCNi-VTA neurons were labeled with GFP, regions containing both TH-positive and GFP-positive neurons were analyzed.

To quantify GluR2 or VGLUT2 signals within the dorsolateral VTA, positive pixels were identified in z-projection images using the Triangle thresholding algorithm in Fiji, and the percentage of positive area relative to the total dorsolateral VTA image area was calculated.

To quantify VGLUT2 puncta in labeled VTAp–DCN axons or GluR2 puncta in TH-positive VTA neurons, 10 μm-thick 3D images were extracted. A median filter (radius = 2) was applied to reduce noise. VGLUT2 or GluR2 signals and corresponding structural markers were segmented independently using the 3D Object Counter plugin in Fiji. The resulting 3D objects were imported into the 3D Manager and designated as ROIs. Overlap between molecular and structural ROIs was quantified as the percentage of intersecting volume relative to total structural volume.

For analysis of individual VGLUT2 puncta within tdT- or GFP-labeled VTAp–DCN axons, 3D images of VGLUT2 signals restricted to VTAp–DCN axons were generated by masking tdT/GFP-negative regions or extracting overlapping regions using the AND function in the Image Calculator. Individual puncta in the resulting images were segmented using the 3D Object Counter plugin, and object volume and density (per volume of labeled axons) were calculated.

To quantify VGLUT2 signals in the vicinity of GFP-labeled DCNi-VTA somata, z-projection images generated from 2.5 μm optical sections were analyzed. Median filtering (radius = 2) and Triangle thresholding were applied. Somatic ROIs were manually delineated around individual GFP-positive DCNi-VTA somata, the proportion of VGLUT2-positive areas was calculated within the ROIs. TH immunostaining was used to classify each GFP-labeled neuron as TH-positive or TH-negative.

To analyze c-Fos-positive DCN neurons, single-field-of-view z-stack images (295 × 295 × 10 μm; 20 slices at 0.5 μm intervals) were acquired from the DN. Z-projection images of c-Fos and DAPI were generated from the z-stacks. C-Fos-positive areas were identified using the Auto Local Threshold function (MidGrey, radius 300) in Fiji. The areas of individual DCN cells were determined based on DAPI signals. Cells in which more than 40% of the DCN cell area was c-Fos-positive were classified as c-Fos-positive cells. The percentage of c-Fos-positive cells among all DCN cells was then calculated.

### Whole-cell patch-clamp recording

For acute slice preparations, mice were deeply anesthetized with isoflurane and decapitated either without exposure to 14RS or one day after 14RS, at 7–8 weeks of age. The brain was immediately removed, and coronal midbrain slices (150 μm) containing the VTA were prepared using a vibrating microtome (Leica VT1200S) in ice-cold sucrose-based aCSF containing (in mM): 220 sucrose, 2.5 KCl, 1.25 NaH_2_PO_4_, 25 NaHCO_3_, 20 glucose, 7 MgCl_2_ and 0.5 CaCl_2_, oxygenated with 95% O_2_/5% CO_2_. The slices were incubated for 30 min in warmed (36 °C) normal aCSF, which was the same as that used for western blot sampling. After an additional 1 h incubation at room temperature, slices were transferred to a recording chamber and continuously perfused with oxygenated normal aCSF.

Patch pipettes (3–5 MΩ) were filled with an internal solution containing (in mM): 130 potassium gluconate, 2 NaCl, 4 MgCl_2_, 4 Na_2_-ATP, 0.4 Na-GTP, 20 HEPES (pH 7.2), and 0.25 EGTA. For mEPSC recording, the extracellular solution contained tetrodotoxin (0.5 μM; Tocris Bioscience) and bicuculline (1 μM; Tocris Bioscience). Whole-cell recordings were performed on VTA neurons visualized under a microscope (Olympus BX61WI). mEPSCs were recorded from DAergic neurons in the dorsolateral VTA at a holding potential of −70 mV. In DAT-IRES-Cre;Ai6 mice, DAergic neurons were identified by ZsGreen fluorescence using an Olympus confocal microscope. However, the injection of rAAV2-retro-CAG-Cre into the VTA of DAT-IRES-Cre;Ai6 mice may induce ZsGreen expression in non-DAergic neurons. Therefore, in experiments involving chemogenetic inhibition or VGLUT2 OE in VTAp–DCN neurons, Ih currents were measured by applying hyperpolarizing voltage steps (−110 mV) from a holding potential of −60 mV. The amplitude of Ih currents was calculated as the difference between the maximum and minimum values during 500 ms of voltage steps, as described previously ^10^. Neurons exhibiting Ih currents larger than 15 pA were classified as DAergic neurons in both DAT-IRES-Cre;Ai6 and C57BL/6J mice. Recordings were performed using a MultiClamp 700B amplifier, and data were acquired with pCLAMP software (Molecular Devices). Recordings were included for analysis if input membrane resistance exceeded 200 MΩ, and the holding current was less than 200 pA. Cells showing no mEPSC events were excluded from amplitude analysis. Recorded traces were filtered using Clampfit (Molecular Devices), and mEPSCs were detected and analyzed using the Peak Analyzer function in OriginPro (OriginLab). To summarize the recordings, mEPSC frequency, inter-event interval, and peak amplitude were measured.

### Fiber photometry measurements

To measure DA release, a fiber-optic cannula was implanted 2 weeks after injection of AAV-hSyn-GRAB_DA2h_ into the NAc. A cannula with a 1.25 mm ceramic ferrule (400 μm core diameter, 0.39 NA, 4 mm length; RWD Life Science) was inserted 0.2 mm above the AAV injection site and secured with dental cement (Superbond C&B, Sun Medical) and self-curing resin (Vertex). After approximately 5 days of recovery, mice underwent the first fiber photometry recording, followed by a second recording 2 weeks later, either with or without 14RS during the interval.

A low-autofluorescence fiber-optic patch cord (Doric Lenses Inc.) was connected to the implanted cannula. A 465 nm LED (modulated at 330 Hz) was used to excite GRAB_DA2h_, and a 405 nm LED (modulated at 210 Hz) was used as an isosbestic control for motion artifacts and photobleaching. Light from these LEDs was delivered through a filtered minicube (ilFMC4, Doric Lenses Inc.). Signals were processed using a fiber photometry processer (RZ5P, Tucker Davis Technologies (TDT)) and collected using Synapse software (TDT). Recordings were conducted in the home cage without a lid. LED power was adjusted to 20–30 μW (465 nm) and 2–10 μW (405 nm) to avoid signal saturation. Baseline signals were recorded for 6 min before presentation of a sweetened pellet. The first 3 min were used to assess tonic DA release. Mice were food-deprived for 24 h before both recording sessions. Experiments were conducted between 14:00 and 18:00.

For analysis, custom Python scripts based on the TDT Fiber photometry User Guide (https://www.tdt.com/docs/fiber-photometry-user-guide/data-analysis/) were used. Signals from both 465 nm and 405 nm channels were downsampled (factor = 10). The 405 nm signal was linearly fitted to the 465 nm signal to correct for motion artifacts and photobleaching. Normalized fluorescence was calculated as: ΔF/F_0_ = (465 nm signal – scaled 405 nm signal)/ scaled 405 nm signal.

Z-scores were calculated using baseline values from −330 to −30 s before pellet biting (defined as t = 0). For z-score traces, dark lines represent the mean and shaded areas represent SEM, with results from the two recordings overlaid. Peak feeding-evoked phasic DA responses were detected within 0–30 s after pellet biting, given that sustained signals tended to decrease during the second recording even in CTR mice, as shown by the time-dependent changes. Peak responses from the first and second recordings were normalized to the first recording in each mouse and compared between recordings. For group comparisons, the ratio of the second to the first peak response (R_DA-phasic_) was calculated. Because recordings were performed twice in the same mice, changes in tonic DA release were estimated by comparing baseline GRAB_DA2h_ signals between recordings. The average 465 nm signal divided by the average 405 nm signal during the first 3 min was calculated, and the ratio of the second to the first measurement (R_DA-tonic_) was derived.

### Fluorescence in situ hybridization (FISH)

C57BL/6J mice with or without 14RS were perfused as described above. Brains were post-fixed in 4% PFA for 24 h at 4°C and cryoprotected in 10%, 20%, and 30% sucrose in PBS until they sank. Coronal sections (20 μm) were prepared using a cryostat (Leica CM1800). Sections were briefly rinsed in PBS to remove optimal cutting temperature compound, mounted onto Superfrost glass slides (Fisher Scientific), and stored at –80°C until use. FISH was performed using the RNAscope Multiplex Fluorescent Reagent Kit v2 (ACDbio), following the manufacturer’s instructions, except that the target retrieval step was omitted to minimize tissue damage. Slc17a6 (VGLUT2) mRNA was detected using probe 319171-C2 (ACD) and visualized with Opal 570 (1:500 dilution; Akoya Biosciences).

Single-field-of-view z-stack images (295 × 295 × 5 μm, 10 slices at 0.5 μm intervals) were acquired using a Nikon AX confocal microscope (60× objective, 1,024 × 1,024 resolution). For quantification of Slc17a6 signals, z-projection images were generated from the z-stacks, and positive areas were detected using the Intermodes thresholding algorithm in Fiji. Slc17a6-positive areas were normalized to the number of Slc17a6-positive neurons per image. Images containing fewer than 10 Slc17a6-positive neurons were excluded from analysis to minimize variability due to small denominators.

### Statistical analysis

Statistical analyses were performed using Origin Pro software. Data are presented as boxplots with dots representing individual data points, center lines denoting the median, the lower and upper bounds of the box corresponding to the 25^th^ and 75^th^ percentiles, respectively, and whiskers indicating the most extreme data points within 1.5× the interquartile range from the edges of the box. One-way ANOVA followed by Fisher’s least significant difference (LSD) post hoc test was used for comparisons among more than two groups. For two-group comparisons, the Mann–Whitney U test was used for sample sizes smaller than 10 (n ≤ 10), and a two-tailed unpaired Student’s t-test was used when n > 10. Paired two-sample t-tests were used for within-mouse comparisons of fiber photometry recordings. A p value of less than 0.05 was considered to indicate a statistically significant difference between groups. All statistical results are summarized in Table S1, and the sample numbers in individual groups are presented both in the figure legends and Table S1.

## Supplemental figure legend

**Figure S1.**
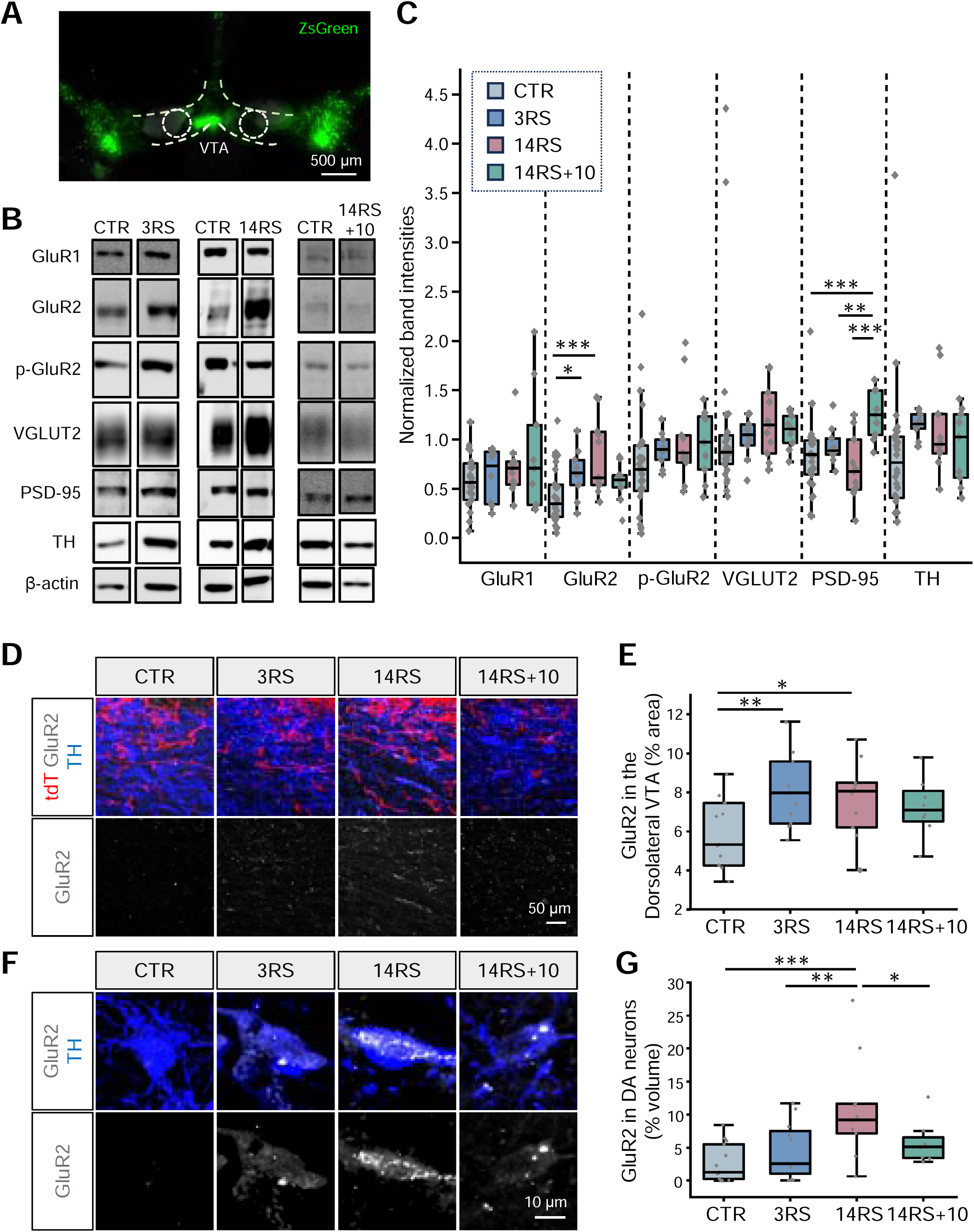
RS-dependent changes in excitatory synaptic molecules (Related to Figure 1) (A) Representative image of a midbrain slice from a DAT-IRES-Cre;Ai6 mouse after bilateral VTA regions were collected using a small punch. (B) Representative immunoblots probed with antibodies against GluR1, GluR2, p-GluR2, VGLUT2, PSD-95, TH, and β-actin in VTA lysates. Each 3RS, 14RS, or 14RS+10 group was compared with CTR. (C) Quantification of immunoblots, with band intensities normalized to β-actin in each sample. Sample sizes were CTR: n = 26 mice; 3RS: n = 9 mice; 14RS: n = 11 mice; 14RS+10: n = 9 mice, except for GluR1 (CTR: n = 22 mice; 3RS: n = 8 mice; 14RS: n = 10 mice; 14RS+10: n = 9 mice). (D) Representative 3D projection images of the dorsolateral VTA stained for TH (blue) and GluR2 (gray), with tdT-labeled DCN axons (red), in CTR, 3RS, 14RS, and 14RS+10 groups. (E) Percentage of GluR2-positive area in the dorsolateral VTA (CTR: n = 11 mice; 3RS: n = 11 mice; 14RS: n = 10 mice: 14RS+10: n = 8 mice). (F) Representative 3D projection images of GluR2 staining (gray) restricted to TH-positive DAergic neurons (blue) in the dorsolateral VTA of CTR, 3RS, 14RS, and 14RS+10 groups. (G) Percentage of GluR2-positive volume within TH-positive DAergic neurons (CTR: n = 13 mice; 3RS: n = 12 mice; 14RS: n = 11 mice; 14RS+10: n = 8 mice). *p < 0.05, **p < 0.01, *** < 0.001; one-way ANOVA followed by Fisher’s LSD post hoc test.

**Figure S2.**
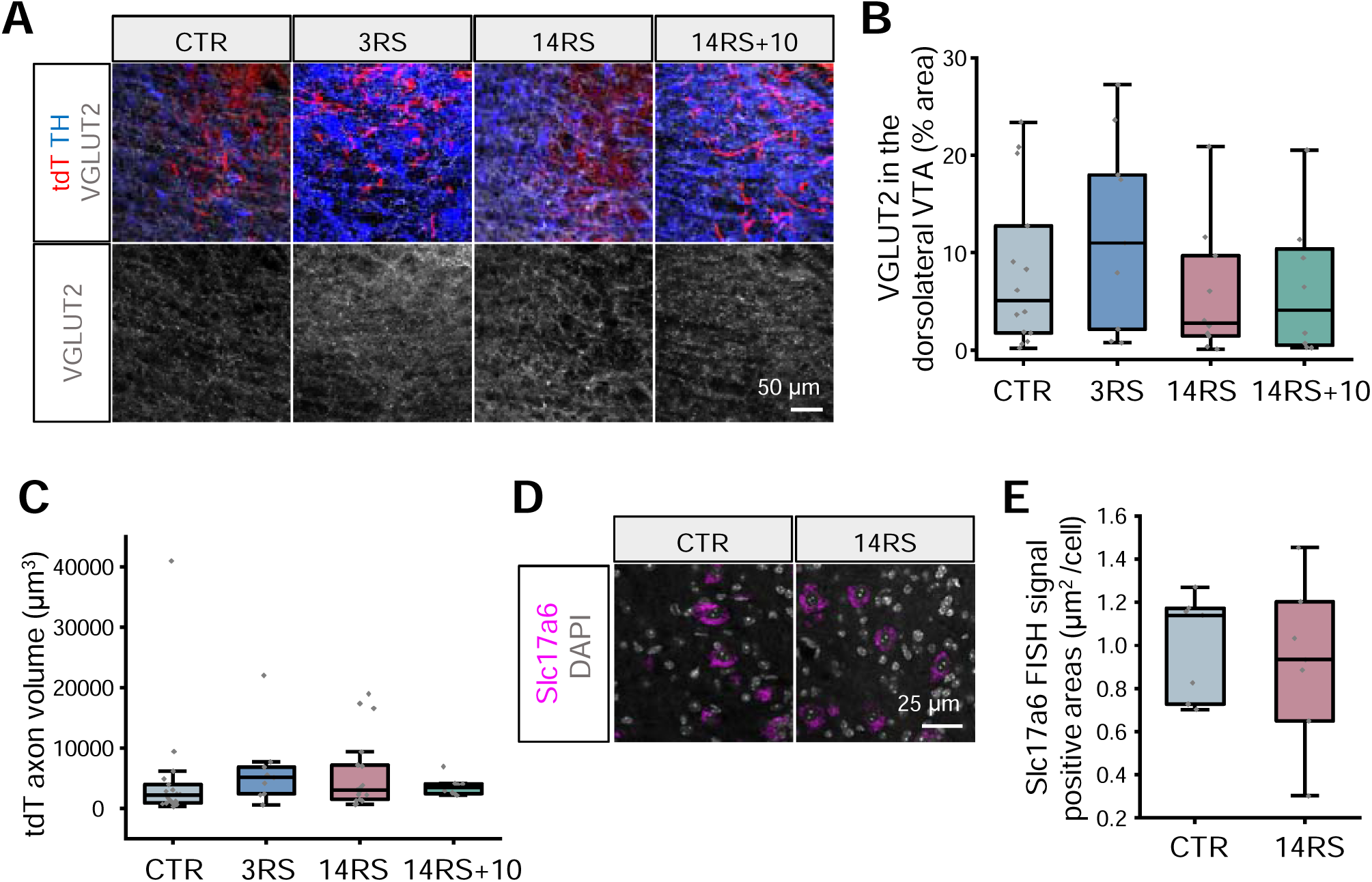
Unaltered overall VGLUT2 expression, VTAp–DCN axonal volume, and VGLUT2 mRNA expression (Related to Figure 1) (A) Representative 3D projection images of the dorsolateral VTA stained for TH (blue) and VGLUT2 (gray), with tdT-labeled DCN axons (red), in CTR, 3RS, 14RS, and 14RS+10 groups. (B) Percentage of VGLUT2-positive area in the dorsolateral VTA in CTR (n = 14 mice), 3RS (n = 9 mice), 14RS (n = 10 mice), and 14RS+10 (n = 8 mice). (C) Volume of tdT-positive VTAp–DCN axons per image in the dorsolateral VTA (CTR: n = 19 mice; 3RS: n = 9 mice; 14RS: n = 17 mice; 14RS+10: n = 18 mice). (D) Representative 3D projection images of Slc17a6 mRNA (magenta) and DAPI (gray) in the DN of CTR and 14RS mice. (E) Average Slc17a6 FISH signal-positive area per DCN neuron (n = 7 mice per group). No significant differences were detected by one-way ANOVA (B and C) or Mann–Whitney U test (E).

**Figure S3.**
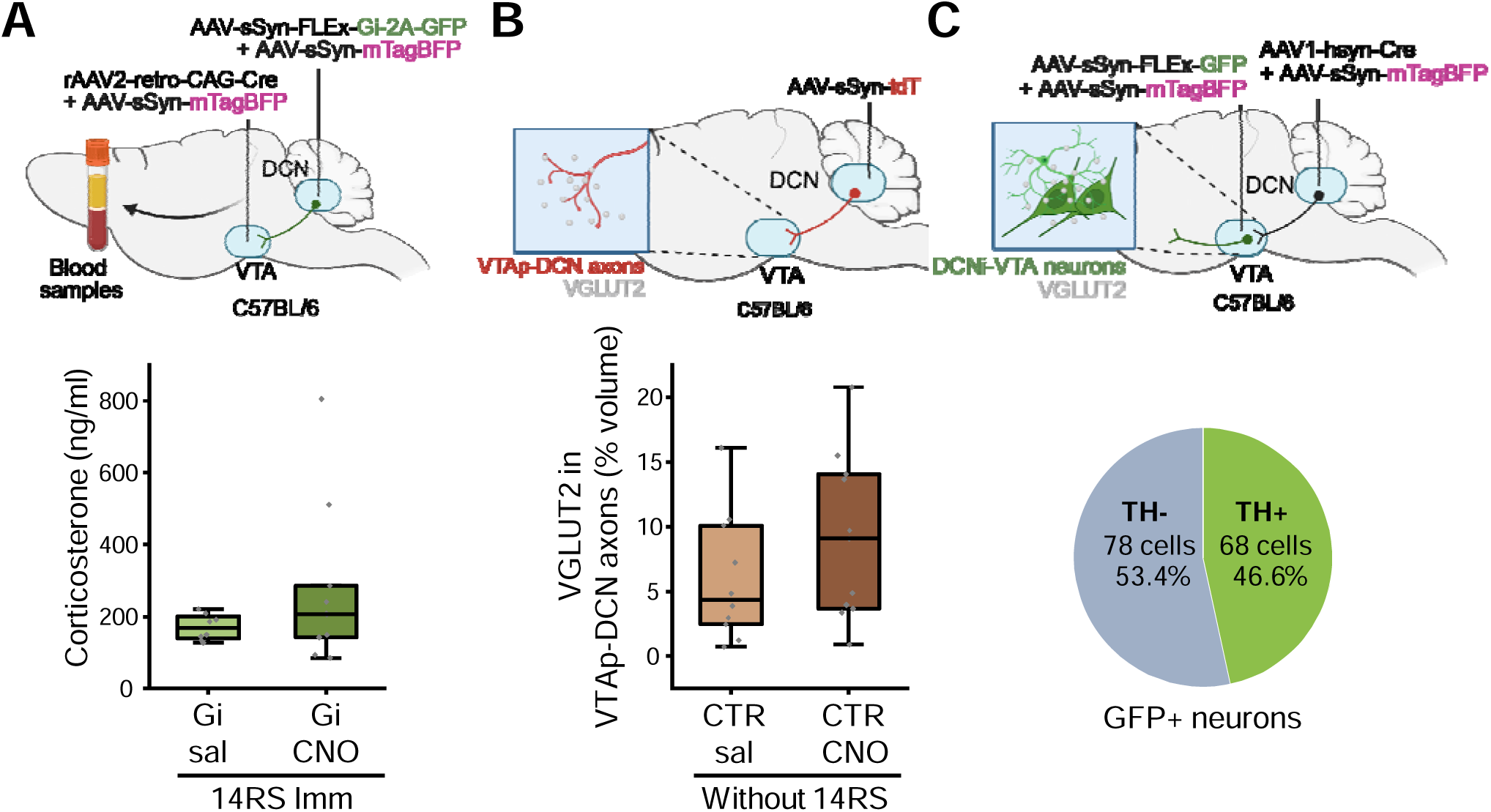
Control experiments confirming activity-dependent reduction of VGLUT2 in VTAp–DCN axons (Related to Figure 2) (A) Serum corticosterone levels measured immediately after 14RS (14RS Imm) in mice expressing Gi in VTAp–DCN neurons following 14 days of saline (Gi-sal; n = 8 mice) or CNO (Gi-CNO; n = 10 mice) administration. (B) Percentage of VGLUT2-positive volume within tdT-labeled VTAp–DCN axons in CTR mice following 14 days of saline (CTR-sal; n = 10 slices from 6 mice) or CNO (CTR-CNO; n = 11 slices from 6 mice) administration. (C) Pie chart illustrating proportions of TH-positive (n = 63) and TH-negative (n = 80) neurons among GFP-labeled DCNi-VTA neurons. No significant differences were detected by Mann–Whitney U test (A and B).

**Figure S4.**
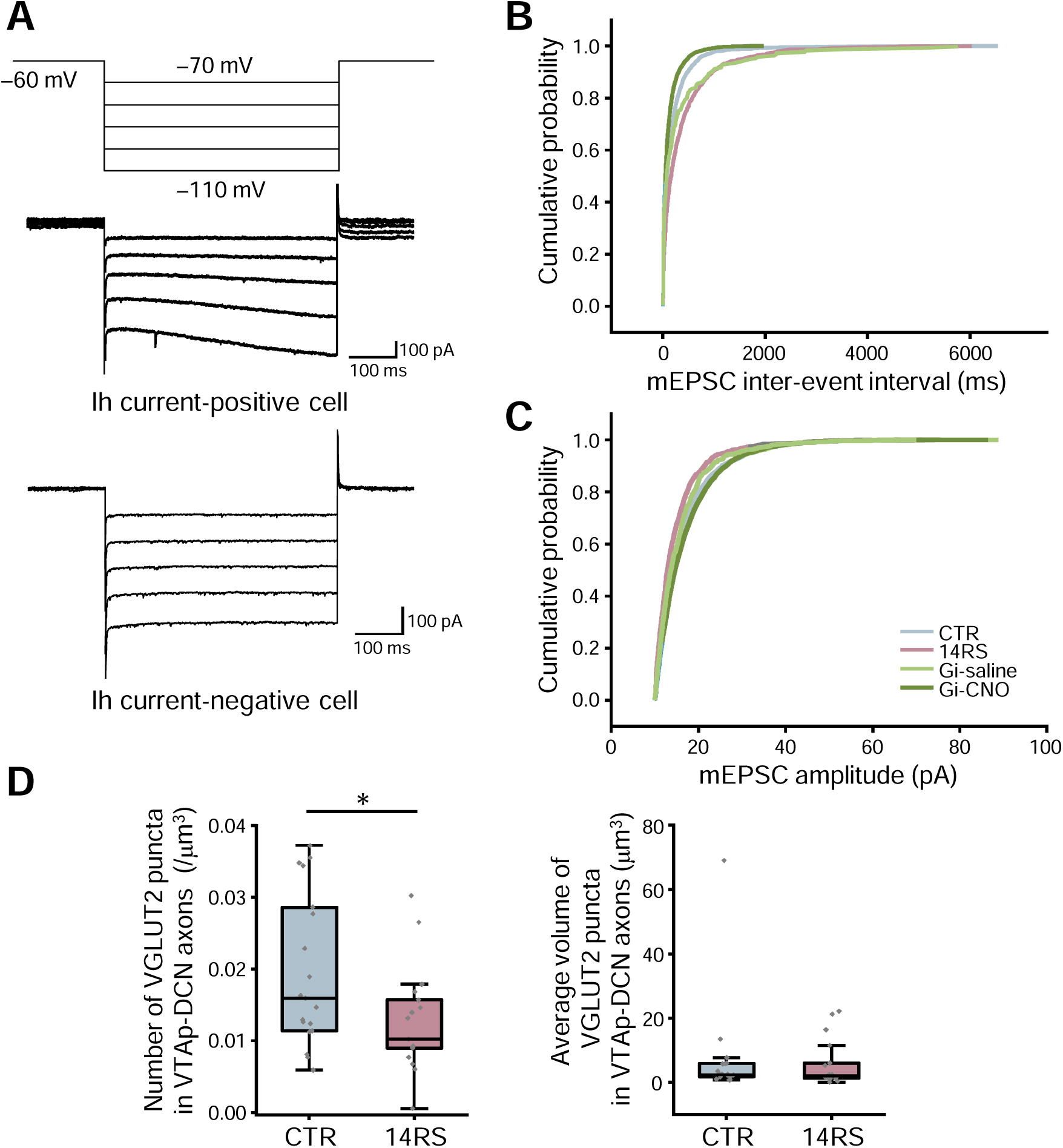
Potential link between reduced VGLUT2 puncta number and decreased mEPSC frequency (Related to Figure 3) (A) Representative patch-clamp recordings showing currents evoked by voltage steps from −70 mV to −110 mV (holding potential: −60 mV). (B and C) Cumulative probability distributions of mEPSC inter-event intervals (B) and amplitude (C) in CTR (n = 9 cells from 5 mice), 14RS (n = 11 cells from 11 mice for (B); n = 10 cells from 10 mice for (C)), Gi-sal (n = 6 cells from 4 mice for (B); n = 5 cells from 3 mice for (C)), and Gi-CNO (n = 7 cells from 4 mice) groups. (D) Number (left) and average volume (right) of VGLUT2-positive puncta in VTAp–DCN axons in CTR (n = 19 mice) and 14RS (n = 17 mice) groups. *p < 0.05; two-tailed unpaired Student’s t-test (D).

**Figure S5.**
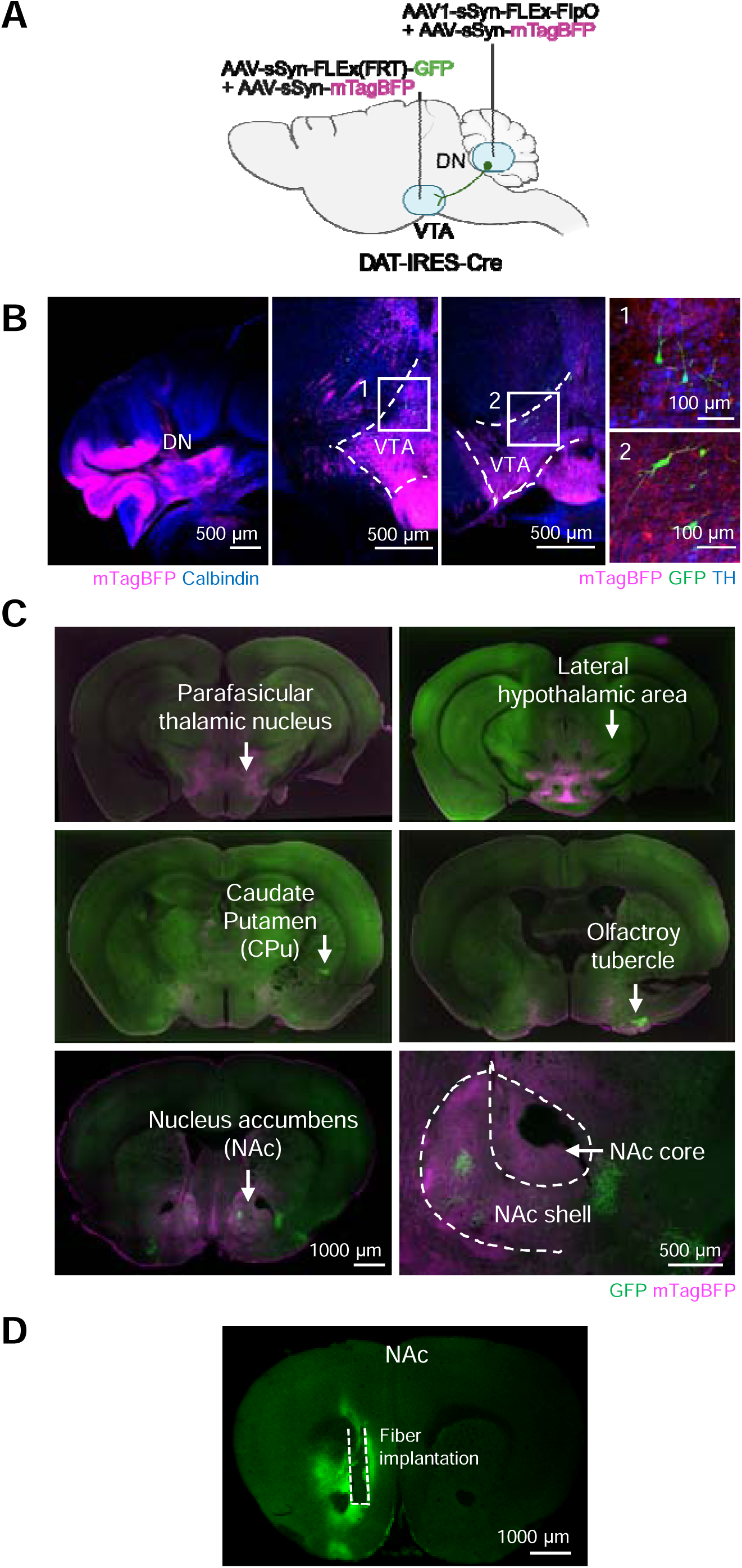
Projections of DAergic DCNi-VTA neurons (Related to Figure 3) (A) Schematic of AAV injection for labeling DAergic DCNi–VTA neurons. (B) Representative images of mTagBFP expression in the cerebellum and GFP/mTagBFP expression in the VTA. Regions within white squares in the dorsolateral VTA are magnified on the right. (C) GFP-positive axonal projections arising from DAergic DCNi-VTA neurons across multiple brain regions, including dense projections to the NAc. The NAc region is shown at higher magnification in the lower right. (D) Representative midbrain slice illustrating the fiber implantation site and GRAB_DA2h_ expression for fiber photometry measurements.

**Figure S6.**
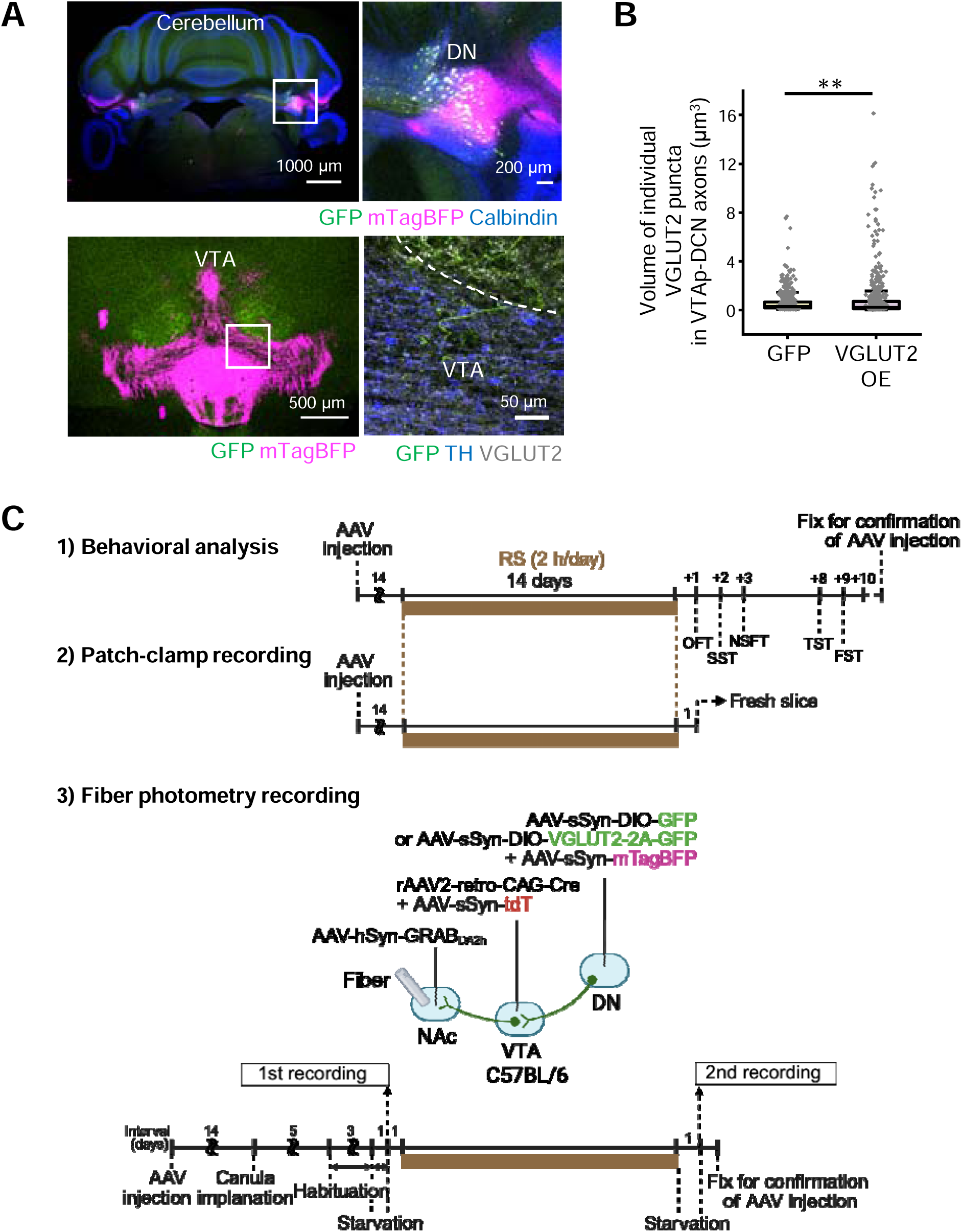
Effects of VGLUT2 OE in VTAp–DCN neurons (Related to Figure 4) (A) Representative images of GFP and mTagBFP expression in the DCN and VTA following AAV injection shown in Figure 4A. Regions outlined by white squares are shown at higher magnification on the right. (B) Individual VGLUT2 puncta volume within GFP-positive VTAp–DCN axons after GFP expression alone (n = 528 puncta from 5 mice) or VGLUT2 OE (n = 614 puncta from 6 mice). The average volume per mouse derived from the same data is shown in Figure 4C. (C) Timelines for behavioral tests, patch-clamp recordings, and fiber photometry experiments following AAV-mediated VGLUT2 OE in VTAp–DCN axons. **p < 0.01; two-tailed unpaired Student’s t-test.

**Table S1 (Excel file).** P values, statistical tests, other statistical parameters, and sample sizes related to Figures 1–5 and S1–S6.

## Key resources table

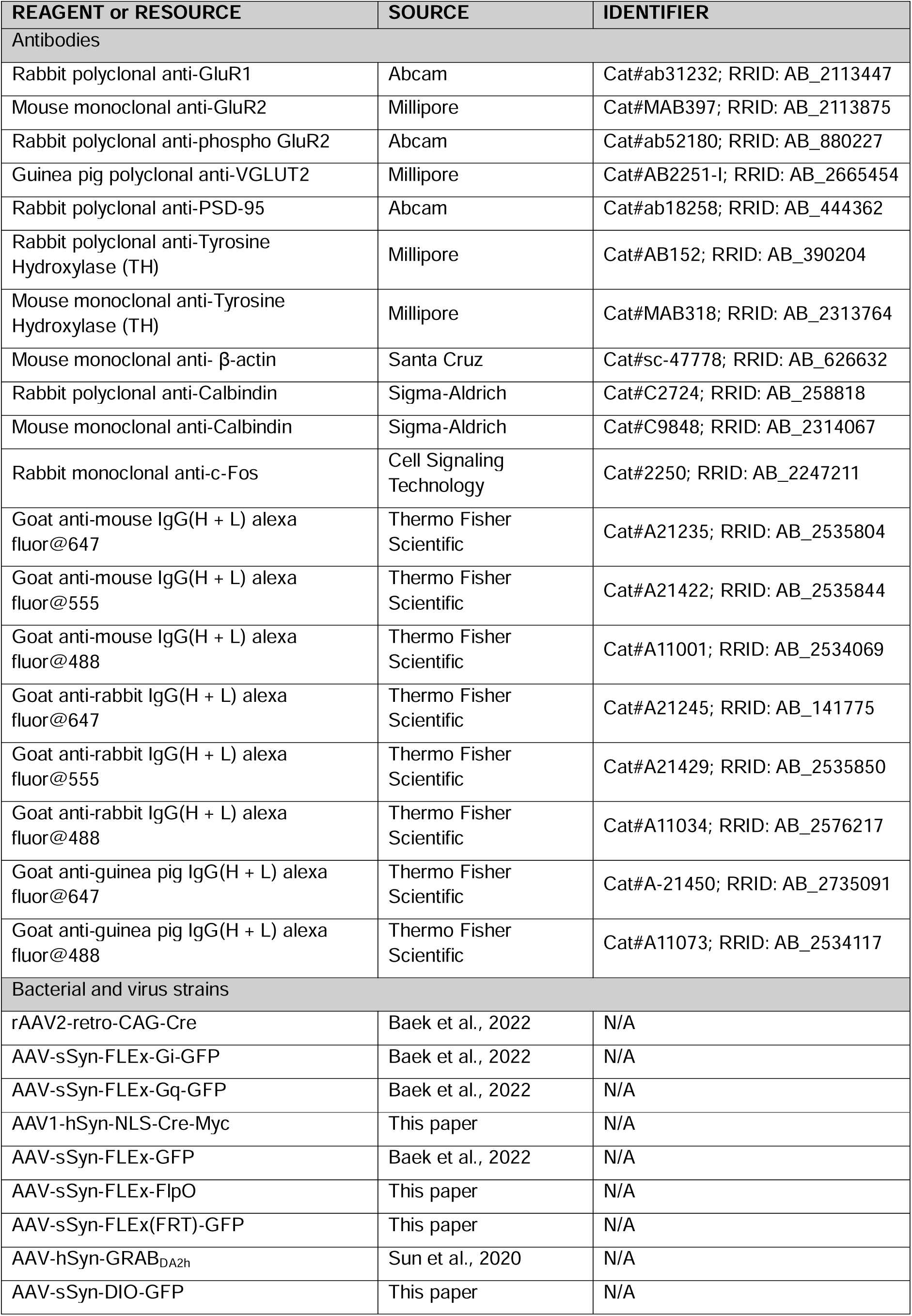
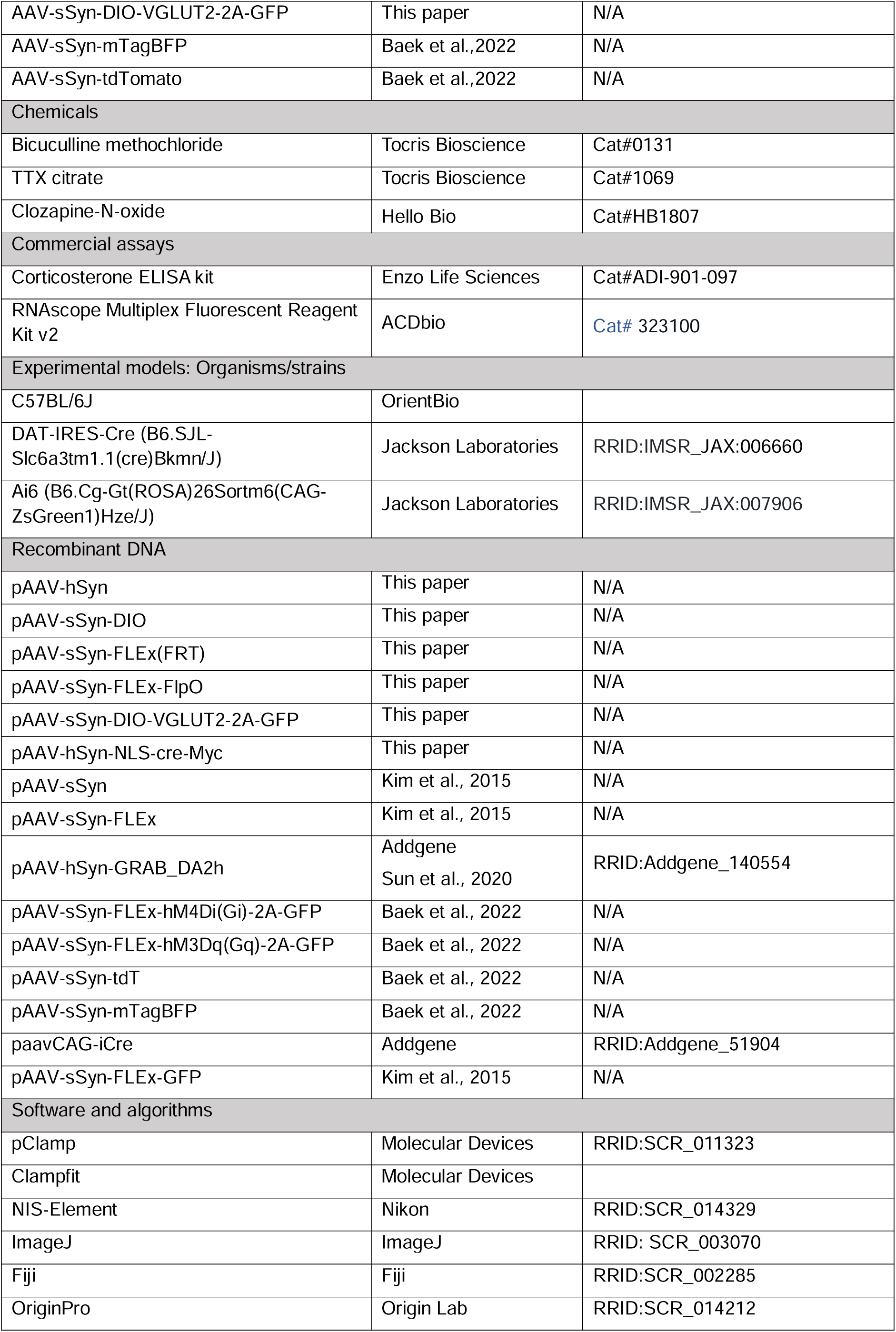
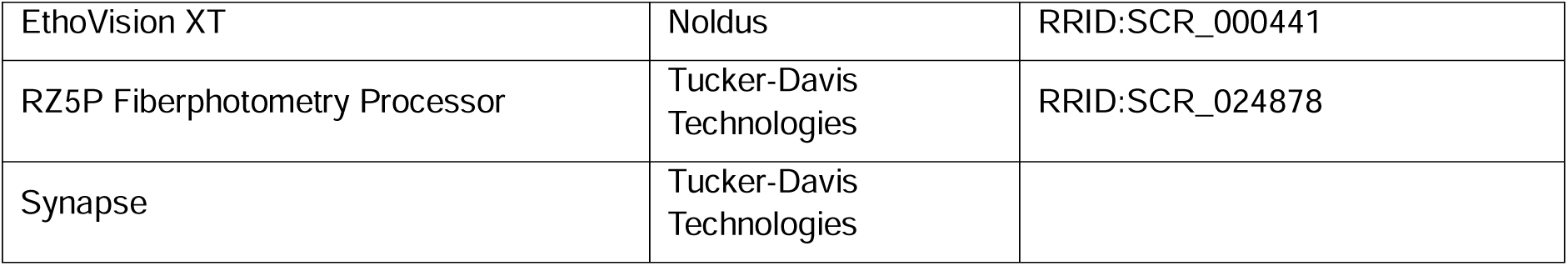

## Notes

### Competing Interest Statement

The authors have declared no competing interest.

